# Dietary lipid stress aggravates obstructive lung injury involving epithelial IGF-1-Akt dysfunction and epithelial-endothelial crosstalk

**DOI:** 10.64898/2026.01.29.702619

**Authors:** Choyo Ogasawara, Hirofumi Nohara, Tomoki Kishimoto, Keisuke Kawano, Kenji Watanabe, Kentaro Oniki, Yukio Fujiwara, Ryunosuke Nakashima, Shunsuke Kamei, Noriki Takahashi, Megumi Hayashi, Ayami Fukuyama, Mai Uemura, Keiko Ueno-Shuto, Junji Saruwatari, Yasuhiro Ogata, Yoichi Mizukami, Mary Ann Suico, Hirofumi Kai, Tsuyoshi Shuto

**Affiliations:** Department of Molecular Medicine, Graduate School of Pharmaceutical Sciences, Kumamoto University, 5-1 Oe-Honmachi, Chuo-ku, Kumamoto 862-0973, Japan; Health Life Science S-HIGO Professional Fellowship Program, Kumamoto University, 2-39-1 Kurokami, Chuo-ku, Kumamoto 862-8555, Japan; Program for Fostering Innovators to Lead a Better Co-being Society, Kumamoto University, 2-39-1 Kurokami, Chuo-ku, Kumamoto 862-8555, Japan; Center for Gene Research, Science Research Center, Yamaguchi University, Ube, Yamaguchi 755 8505, Japan; Division of Pharmacology and Therapeutics, Graduate School of Pharmaceutical Sciences, Kumamoto University, 5-1 Oe-Honmachi, Chuo-ku, Kumamoto 862-0973, Japan; Department of Cell Pathology, Graduate School of Medical Science, Kumamoto University, 1-1-1 Honjo, Chuo-ku, Kumamoto, 860-8556, Japan; Laboratory of Pharmacology, Division of Life Science, Faculty of Pharmaceutical Sciences, Sojo University, 4-22-1 Ikeda, Nishi-ku, Kumamoto 860-0082, Japan; Japanese Red Cross Kumamoto Health Care Center, Kumamoto, 2-1-1 Nagamine-minami, Higashi-ku, Kumamoto, 861-8520, Japan; Global Center for Natural Resources Sciences, Faculty of Life Sciences, Kumamoto University, 5-1 Oe-Honmachi, Chuo-ku, Kumamoto, 862-0973, Japan

**Keywords:** COPD, lipotoxicity, high-fat diet, airway epithelium, IGF-1, Akt, endothelium, epithelial-endothelial crosstalk, hepatic steatosis

## Abstract

**Background:** Altered lipid metabolism is increasingly implicated in chronic obstructive pulmonary disease (COPD), but it remains unclear how a pre-existing obstructive lung state modifies the response to systemic lipid excess. We investigated whether high-fat diet (HFD) amplifies COPD-relevant lung injury and examined epithelial and vascular programs associated with this response.

**Methods:** Male wild-type (WT) and beta-epithelial sodium channel-transgenic (βENaC-Tg) mice were fed control diet or HFD for 10-11 weeks. Lung structure and function, whole-lung transcriptomes, Akt-FOXO1 signaling, apoptosis-related responses, and pulmonary vascular profiles were assessed. Streptozotocin-induced insulin-deficient diabetes and pharmacological IGF-1 receptor inhibition were used as mechanistic comparators. Palmitate responses were examined in human bronchial epithelial cells, ENaC-hyperactive epithelial cells, and endothelial cells, including conditioned-medium transfer. HFD preconditioning was also evaluated in an elastase-induced emphysema model. Statistical analyses included unpaired two-tailed Student’s *t* tests, one-way ANOVA with Tukey-Kramer or Dunnett multiple-comparison testing, Pearson correlation, and Benjamini-Hochberg correction for RNA-sequencing analyses.

**Results:** HFD produced similar increases in body weight, glycemia, and adiposity in WT and βENaC-Tg mice, while further increasing distal-airspace enlargement and reducing FEV0.1/FVC in βENaC-Tg mice. Lung transcriptomics revealed coordinated remodeling of lipid metabolic, PI3K-Akt, and vascular programs. HFD reduced Akt phosphorylation, increased FOXO1 and *Fasl*, and increased TUNEL-positive cells in epithelial regions. Palmitate attenuated IGF-1-induced Akt activation in bronchial epithelial cells, whereas IGF-1 receptor inhibition reproduced Akt suppression and apoptosis-related responses without fully reproducing the HFD phenotype. HFD preconditioning also increased elastase-induced airspace enlargement, accompanied by parallel upregulation of FOXO1 and TUNEL positivity. HFD reduced pulmonary CD34-positive vascular profiles, and palmitate activated endothelial cells directly and through conditioned media from ENaC-hyperactive epithelium. In men with airflow obstruction, hepatic steatosis coincided with lower percent-predicted FEV1.

**Conclusions:** Dietary lipid stress amplifies obstructive lung injury and engages complementary epithelial and vascular responses. Impaired epithelial IGF-1-Akt signaling and epithelial-endothelial crosstalk provide a mechanistic framework linking systemic metabolic stress to reduced resilience of the obstructive lung.

## Background

Chronic obstructive pulmonary disease (COPD) is a heterogeneous lung condition characterized by chronic respiratory symptoms and persistent airflow obstruction arising from abnormalities of the airways and/or alveoli [1]. COPD remains a major contributor to the global burden of chronic respiratory disease [2]. Its clinical course is strongly influenced by extrapulmonary comorbidities, including cardiovascular and metabolic disorders, which shape disease heterogeneity, symptom burden, and prognosis [3,4]. Obesity is common in COPD, although its respiratory consequences vary with body composition, fat distribution, disease severity, and the accompanying metabolic phenotype [5]. Contemporary treatable-trait frameworks therefore recognize systemic metabolic features as potentially actionable modifiers of COPD biology [6].

Lipids are indispensable for pulmonary physiology, yet excess or maladaptive lipid handling can disrupt tissue homeostasis. Lipotoxicity arises when cellular lipid exposure exceeds the capacity for safe storage, oxidation, or membrane remodeling [7]. Pulmonary surfactant is intrinsically lipid rich, and alveolar epithelial cells require tightly regulated lipid synthesis and transport to preserve alveolar stability [8]. Alveolar type II epithelial cell fatty-acid synthase supports surfactant lipid homeostasis and limits cigarette-smoke-induced experimental COPD [9], while airway epithelial repair requires metabolic rewiring toward fatty-acid oxidation [10]. Thus, the pulmonary effects of lipids depend on the lipid species, cellular context, and the tissue’s capacity to adapt.

Multiple lipid classes have been linked to COPD pathogenesis. Bioactive sphingolipids, including ceramides, participate in apoptosis, oxidative stress, inflammation, and emphysema [11,12], and circulating sphingolipid profiles distinguish COPD phenotypes [13]. Altered glycerophospholipid metabolism has also been detected in human COPD lung tissue [14]. Recent reviews and plasma lipidomic studies further describe broad lipid-metabolic reprogramming and disease-specific lipid signatures in COPD [15,16].

Mechanistic studies have begun to define how systemic lipid dysregulation injures the lung. Severe diet-induced obesity can generate COPD-like pathology through macrophage PLA2G7-dependent accumulation of arachidonic acid, oxidative lipid stress, NLRP3 inflammasome activation, and pyroptosis [17]. Excess cholesterol can aggravate cigarette-smoke-induced injury through macrophage mitochondrial dysfunction and ROS-dependent PPIA-NF-κB signaling [18]. Lipid-associated communication between lung compartments is also emerging: macrophage-derived extracellular vesicles containing ceramides impair endothelial homeostasis [19], and extracellular vesicles from COPD airway epithelial cells can induce endothelial inflammation and apoptosis [20]. These findings establish immune and vascular pathways of lipid-associated injury, while leaving the metabolic response of the diseased airway epithelial-vascular unit incompletely defined.

We reasoned that the pre-existing state of the lung may determine how systemic lipid excess is translated into tissue injury. Chronic mucus obstruction and epithelial stress could reduce the adaptive reserve of the airway, allowing a metabolic challenge that is tolerated by a relatively normal lung to amplify established disease. The phosphatidylinositol-3-kinase (PI3K)-Akt pathway is a plausible interface between metabolic exposure and structural-cell survival. Akt inhibits FOXO transcription factors and BAD [21,22], regulates endothelial signaling and vascular homeostasis [23,24], and is activated by insulin-like growth factor-1 (IGF-1), a growth factor with established roles in lung development and repair [25].

To test this concept, we first examined HFD in βENaC-Tg mice, which develop chronic airway-surface dehydration, mucus obstruction, epithelial stress, inflammation, and distal-airspace abnormalities [26–28]. We combined physiological analysis with whole-lung transcriptomics and mechanistic experiments focused on epithelial IGF-1-Akt signaling and epithelial-endothelial communication. We then assessed whether HFD preconditioning also increased susceptibility to a subsequent elastase-induced emphysema-like insult [28,29]. Finally, we examined hepatic steatosis and lung function in a health-screening cohort to place the experimental findings in a human metabolic context.

## Methods

### Study design

This study integrated two COPD-relevant mouse models, whole-lung transcriptomics, epithelial and endothelial cell systems, pharmacological pathway analysis, and a retrospective clinical cohort. The primary experiment compared WT and βENaC-Tg mice maintained on control diet or HFD. Complementary experiments evaluated HFD exposure before elastase-induced emphysema, short-term streptozotocin-induced insulin-deficient diabetes, IGF-1 receptor inhibition, direct palmitate exposure, and bidirectional epithelial-endothelial conditioned-medium transfer. The biological experimental unit was one mouse for animal analyses and one independently performed cell culture experiment for *in vitro* analyses. Multiple microscopic fields, wells, or technical assay spots from the same biological unit were averaged before statistical analysis. The Akt antibody array was performed once per condition and was used only for exploratory visualization; duplicate antibody spots were technical replicates.

### Animals and ethical approval

Male C57BL/6J WT mice were obtained from Charles River Japan. βENaC-Tg mice were generated and maintained at the Center for Animal Resources and Development, Kumamoto University, as previously described [26,27]. The colony was derived from B6.Cg-Tg(Scgb1a1-Scnn1b)6608Bouc/J mice (JAX stock no. 006438; RRID: IMSR_JAX:006438). Genotyping was performed according to the established colony protocol [27]. Mice were maintained under specific-pathogen-free conditions on a 12-h light/12-h dark cycle at 22-25°C with ad libitum access to water and diet. All animal experiments were approved by the Animal Welfare Committee of Kumamoto University (approval no. A2024-052) and were conducted in accordance with institutional and national guidelines and ARRIVE recommendations. For terminal procedures, mice were anesthetized by intraperitoneal administration of medetomidine (0.75 mg/kg; Nippon Zenyaku Kogyo Co., Fukushima, Japan; Cat. No. 14111), midazolam (4 mg/kg; Sandoz Canada Inc., Canada; Cat. No. 1124401A1060), and butorphanol (5 mg/kg; Meiji Seika Pharma Co., Ltd., Japan; Cat. No. 817000470). After confirmation of adequate anesthesia, blood was collected by cardiac puncture and mice were euthanized by exsanguination.

### Dietary intervention and metabolic assessment

Mice were maintained on standard control diet (MF laboratory chow; Oriental Yeast Co., Tokyo, Japan) or a custom HFD supplied by Oriental Yeast Co. for 10-11 weeks beginning at 5-6 weeks of age unless otherwise specified. The principal components of the HFD included 25% casein, 20% safflower oil, 14% beef tallow, 8% dextrin, 7% lactose, 7% sucrose, and 5% cellulose, as used in our previous metabolic study [30]. Body weight was recorded weekly. Fasting and non-fasting blood glucose concentrations were measured using an Accu-Chek Compact glucometer (Roche, USA). At the endpoint, subcutaneous and epididymal adipose depots were excised and weighed. For oral glucose-tolerance testing, mice were fasted for 6 h and administered glucose (1 g/kg) by oral gavage; blood glucose was measured at 0, 15, 30, and 60 min.

### βENaC-Tg muco-obstructive lung disease model

WT and βENaC-Tg mice were maintained on control diet or HFD for 10-11 weeks. The βENaC-Tg model was used to represent a chronic muco-obstructive disease background characterized by increased epithelial Na^+^ absorption, airway-surface dehydration, mucus obstruction, inflammation, and distal-airspace abnormalities [26–28]. Metabolic measurements, pulmonary function, lung histology, RNA sequencing, and biochemical analyses were performed at the endpoint.

### Elastase-induced emphysema model

Male C57BL/6J mice were maintained on control diet or HFD for 10 weeks and then received porcine pancreatic elastase (60 μg in sterile saline; Sigma-Aldrich, St. Louis, MO, USA; Cat. No. E7885) by intratracheal instillation [28,29]. Akt-FOXO1 signaling was assessed 7 days after elastase administration, whereas airspace enlargement and TUNEL staining were assessed 21 days after elastase administration.

### Streptozotocin-induced insulin-deficient diabetes

Thirteen-week-old male βENaC-Tg mice received a single intraperitoneal injection of STZ (200 mg/kg in sterile saline; Sigma-Aldrich; Cat. No. 572201). Control mice received vehicle. STZ induces pancreatic β-cell injury and insulin-deficient diabetes [31]. Blood glucose was monitored throughout the experiment. Serum insulin was quantified 3 weeks after STZ administration using a mouse insulin ELISA kit (Morinaga Institute of Biological Science, Kanagawa, Japan; Cat. No. M1104). Lung histology and pulmonary function were assessed at the same time point.

### IGF-1 receptor inhibition

Twelve-week-old βENaC-Tg mice received daily intraperitoneal injections of picropodophyllin (PPP; 60 μg/day; Santa Cruz Biotechnology, USA; Cat. No. sc-204008A) for 14 days. Control mice received vehicle. Lung tissue was collected 2 h after the final administration for Akt signaling, histological, apoptotic, and pulmonary function analyses. A corresponding PPP experiment was performed in elastase-treated mice.

### Pulmonary-function measurements

Pulmonary mechanics were measured using a flexiVent small-animal ventilator (SCIREQ Scientific Respiratory Equipment Inc., Canada). Deeply anesthetized mice underwent tracheostomy and tracheal cannulation and were mechanically ventilated at 150 breaths/min as previously described [27]. Inspiratory capacity, static compliance, and elastance were measured using standardized flexiVent perturbations. Forced-expiration maneuvers were used to determine FEV0.1, FVC, and FEV0.1/FVC.

### Lung tissue preparation, histology, and morphometry

After physiological measurements, lungs were perfused with phosphate-buffered saline and fixed in 10% neutral-buffered formalin for 12 h before dehydration and paraffin embedding. Sections were cut at 6 μm and stained with H&E, PAS, or Alcian blue as appropriate. Hematoxylin solution Gill No. 3 (Sigma-Aldrich; Cat. No. GHS332), periodic acid solution (Sigma-Aldrich; Cat. No. 395132), Schiff’s reagent (Sigma-Aldrich; Cat. No. 3952016), and Alcian blue solution (Nacalai Tesque, Kyoto, Japan; Cat. No. 37154-15) were used. Images were acquired using a BZ-X710 microscope (Keyence, Osaka, Japan) under matched settings. Ten non-overlapping fields per lung section were used for alveolar morphometry. MLI, alveolar area, alveolar perimeter, Feret diameter, and mean axis length were quantified using ImageJ (National Institutes of Health, USA). Field-level values were averaged to generate one animal-level value.

### TUNEL analysis

Apoptosis was detected using the DeadEnd Fluorometric TUNEL System (Promega, Madison, WI, USA; Cat. No. G3250) according to the manufacturer’s instructions. Sections were counterstained with DAPI (1 μg/mL; Dojindo Laboratories, Kumamoto, Japan; Cat. No. D523), mounted using VECTASHIELD Mounting Medium (Vector Laboratories, Burlingame, CA, USA; Cat. No. H-1000), and imaged with a BZ-X710 microscope under matched settings. TUNEL-positive nuclei were quantified in anatomically defined epithelial regions. Multiple fields from each animal were averaged to generate one animal-level value.

### CD34 immunohistochemistry

Paraffin-embedded lung sections were cut at 4 μm, deparaffinized, and subjected to heat-induced antigen retrieval using pH 9.0 antigen-retrieval solution (Nichirei Biosciences, Tokyo, Japan; Cat. No. 715291). Sections were incubated with rabbit anti-CD34 antibody (Abcam, USA; Cat. No. ab81289; RRID: AB_1640331), followed by HRP-conjugated secondary antibody. Immunoreactivity was visualized using DAB substrate (Nichirei Biosciences; Cat. No. 343-00901). Whole-slide images were acquired using a NanoZoomer S20 digital slide scanner (Hamamatsu Photonics, Japan; Cat. No. C16300-01) and analyzed using HALO software (Indica Labs, USA). Identical thresholds were used across groups, and one animal-level summary value was used for analysis.

### Cell lines and culture conditions

Human bronchial epithelial 16HBE14o-cells were provided by Dieter C. Gruenert (RRID: CVCL_0112) and have been described previously [32]. 16HBE14o-β/γENaC cells, which stably overexpress β-and γ-ENaC, were provided by Ray A. Caldwell and were used as an ENaC-hyperactive epithelial model [32]. No dedicated RRID is assigned to this derivative line. HUVECs (HUE102) were obtained from the RIKEN BioResource Research Center (RCB5489) [33].

Epithelial cells were maintained in Eagle’s MEM (FUJIFILM Wako Pure Chemical Corporation, Japan; Cat. No. 1010120) supplemented with 10% fetal bovine serum (HyClone, USA; Cat. No. SH30396.03) and 1% penicillin/streptomycin (FUJIFILM Wako; Cat. No. 168-23191) on dishes coated with fibronectin (Sigma-Aldrich; Cat. No. F2006) and collagen (FUJIFILM Wako; Cat. No. 631-00771). HUVECs were cultured in MEM supplemented with 15% fetal bovine serum (Sigma-Aldrich; Cat. No. F7524), 10 ng/mL recombinant FGF-C (FUJIFILM Wako; Cat. No. 067-06591), 5 μg/mL heparin sodium (FUJIFILM Wako; Cat. No. 085-00134), and 1% penicillin/streptomycin on collagen-coated dishes.

### Preparation of palmitate-containing and control media

Sodium palmitate (palmitic acid sodium salt; Thermo Fisher Scientific, USA; Cat. No. 416700050) was dissolved in 50% (v/v) ethanol at 60 °C in a water bath to prepare a 250 mM stock solution. The stock solution was then diluted 1:50 with a sterile 5% (w/v) bovine serum albumin (BSA) solution prepared in phosphate-buffered saline (PBS), yielding a 5 mM palmitate/BSA working solution. Immediately before cell treatment, the 5 mM working solution was diluted 1:10 with serum-free minimum essential medium (MEM; Sigma-Aldrich, St. Louis, MO, USA) to obtain palmitate-containing medium with a final sodium palmitate concentration of 500 μM. The resulting palmitate-containing medium contained 0.5% (w/v) BSA and 0.1% (v/v) ethanol.

Palmitate-free control medium (CON) was prepared separately by diluting sterile 5% (w/v) BSA in PBS 1:10 with serum-free MEM, yielding a final BSA concentration of 0.5% (w/v). Throughout the cell experiments, CON and PA denote the palmitate-free control medium and the 500 μM palmitate-containing medium, respectively. Cells were exposed to CON or PA medium for the indicated durations.

### IGF-1 stimulation

Human bronchial epithelial 16HBE14o-cells were exposed to palmitate-free control medium (CON) or palmitate-containing medium (PA; 500 μM sodium palmitate) for 12 h and then stimulated for 30 min with recombinant human IGF-1 (50 ng/mL; Thermo Fisher Scientific; Cat. No. PHG0078). Akt and FOXO1 phosphorylation were evaluated by immunoblotting.

### Epithelial-endothelial conditioned-medium experiments

Donor 16HBE14o-, 16HBE14o-β/γENaC, or HUVEC cultures were exposed to palmitate-free control medium (CON) or palmitate-containing medium (PA; 500 μM sodium palmitate) for 3 h. The exposure medium was then completely removed and replaced with fresh palmitate-free medium. Conditioned media were collected 3, 6, and 9 h after medium replacement. Epithelial conditioned media were applied to HUVECs, whereas HUVEC-conditioned media were applied to epithelial cells. Recipient cells were incubated with conditioned medium for 30 min before RNA extraction and quantitative RT-PCR.

### RNA isolation, library preparation, and sequencing

Total RNA from whole-lung tissue was extracted using the Maxwell RSC simplyRNA Kit (Promega; Cat. No. AS1390). RNA integrity was assessed using a TapeStation system (Agilent Technologies, Santa Clara, CA, USA); all sequenced samples had RINe values greater than 8.3. Three independent biological samples per group were analyzed. Poly(A)+ RNA was enriched, libraries were prepared using the NEBNext Ultra II Directional RNA Library Prep Kit (New England Biolabs, Massachusetts, USA; Cat. No. E7760S), and 50-bp paired-end sequencing was performed using an Illumina NovaSeq 6000. Reads were demultiplexed using bcl2fastq, trimmed and filtered, and aligned to mouse GRCm39 release 110 using CLC Genomics Workbench version 23.0.2 (QIAGEN, Venlo, the Netherlands).

### Differential-expression and pathway analyses

Gene-level expression was normalized as transcripts per million (TPM), and log_2_(TPM + 1) values were used for pathway-focused analyses. Pairwise differential expression was assessed using two-tailed Student t tests on log_2_(TPM + 1) values, with Benjamini-Hochberg correction in R version 4.1.1. Differentially expressed genes were defined by an absolute fold change greater than 1.5 and false-discovery rate less than 0.05. Canonical-pathway, upstream-regulator, and interaction-network analyses were performed using Ingenuity Pathway Analysis (QIAGEN).

### Quantitative RT-PCR

Total RNA from cultured cells or mouse lung tissue was extracted using RNAiso Plus (TaKaRa, Tokyo, Japan; Cat. No. 9109) and reverse-transcribed using the PrimeScript RT Reagent Kit (TaKaRa; Cat. No. RR036B). Quantitative PCR was performed using TB Green Premix (TaKaRa; Cat. No. RR820B) on a CFX Connect Real-Time PCR Detection System (Bio-Rad Laboratories, USA). Relative expression was calculated using the 2^−ΔΔCt method. Mouse *Fasl* expression was normalized to mouse *18S rRNA*; human *IL6*, *VCAM1*, and *CGN* were normalized to *GAPDH*. Primer sequences are listed in Additional file 1: Table S1.

### Western blotting

Lung tissues or cultured cells were lysed in RIPA buffer containing 50 mM Tris-HCl (pH 7.5), 150 mM NaCl, 1% NP-40, 1% sodium deoxycholate, 1 mM Na_3_VO_4_, and protease-inhibitor cocktail (Sigma-Aldrich; Cat. No. P8340). Lysates were clarified by centrifugation at 12,000 rpm for 15 min at 4°C, and protein concentrations were determined using BCA reagent (Sigma-Aldrich; Cat. No. B9643). Equal amounts of protein were denatured, separated by SDS-PAGE, and transferred to PVDF membranes (Sigma-Aldrich; Cat. No. IPVH00010). Membranes were blocked in 5% skim milk in TBS-T and incubated with primary antibodies diluted in Can Get Signal Solution 1 (TOYOBO, Japan; Cat. No. NKB-101).

Primary antibodies were rabbit anti-phospho-Akt Thr308 (Cell Signaling Technology; Cat. No. 13038; RRID: AB_2629447; 1:1,000), rabbit anti-Akt (Cat. No. 4691; RRID: AB_915783; 1:1,000), rabbit anti-phospho-FOXO1 Ser256 (Cat. No. 84192; RRID: AB_2800035; 1:1,000), rabbit anti-FOXO1 (Cat. No. 2880; RRID: AB_2106495; 1:1,000), rabbit anti-cleaved caspase-3 (Santa Cruz Biotechnology; Cat. No. sc-22171; RRID: AB_637834; 1:1,000), rabbit anti-gamma-tubulin (Santa Cruz Biotechnology; Cat. No. sc-7396-R; RRID: AB_1120814; 1:1,000), and goat anti-beta-actin (Santa Cruz Biotechnology; Cat. No. sc-1616; RRID: AB_630836; 1:1,000). HRP-conjugated secondary antibodies were goat anti-mouse IgG (Jackson ImmunoResearch; Cat. No. 115-035-003; RRID: AB_10015289; 1:2,000), goat anti-rabbit IgG (Cat. No. 111-035-003; RRID: AB_2313567; 1:2,000), or donkey anti-goat IgG (Cat. No. 705-035-003; RRID: AB_2340390; 1:5,000), diluted in Can Get Signal Solution 2. Signals were developed using SuperSignal chemiluminescent substrate (Thermo Fisher Scientific; Cat. No. 34580), acquired with an LAS-4000 mini system (FUJIFILM; product code 28-9558-13), and quantified using Multi Gauge version 3.1 (RRID: SCR_014299). Uncropped immunoblots are provided in Additional file 2. Each blot lane represents an independent biological sample unless otherwise stated.

### Akt-signaling antibody array

Human bronchial epithelial 16HBE14o-cells were treated with control medium (CON) or palmitate-containing medium (PA; 500 μM sodium palmitate) for 12 h and subsequently stimulated with IGF-1 for 30 min. Cell lysates were analyzed using the PathScan Akt Signaling Antibody Array Kit (Cell Signaling Technology; Cat. No. 9474). Chemiluminescent signals were normalized to the array reference spots, and signals from duplicate antibody spots were averaged for each target. The antibody array was used to profile IGF-1-responsive Akt-pathway nodes under palmitate exposure.

### Serum free-fatty-acid measurement

Serum free fatty acids were quantified using the Free Fatty Acid Quantification Colorimetric Assay (Cell Biolabs, San Diego, CA, USA; Cat. No. STA-618) according to the manufacturer’s instructions.

### Human participants and clinical assessment

The clinical analysis included 474 Japanese men who participated in a health-screening program at the Japanese Red Cross Kumamoto Health Care Center. The cohort comprised 185 never-smokers and 289 ever-smokers. Available variables included age, BMI, fasting blood glucose, serum lipids, liver enzymes, spirometry, smoking history, alcohol intake, diabetes, hypertension, and ultrasonographically assessed hepatic steatosis. Smoking status was obtained by face-to-face interview using a structured questionnaire. No human biospecimens were collected.

Airflow obstruction was defined as pre-bronchodilator FEV1/FVC <0.70. Because post-bronchodilator confirmation and complete clinical diagnostic information were not available for all participants, the term “airflow obstruction” is used rather than “definite COPD”. Hepatic steatosis was assessed by ultrasonography based on diffuse hepatic hyperechogenicity, increased echogenicity relative to the kidney, vascular blurring, and deep attenuation.

### Human research ethics

The human study was conducted in accordance with the Declaration of Helsinki and was approved by the ethics committees of the Japanese Red Cross Kumamoto Health Care Center and the Faculty of Life Sciences, Kumamoto University (Genome no. 169). All participants provided written informed consent.

### Statistical analysis

Data are presented as mean ± SEM unless otherwise specified. Clinical characteristics are presented as mean ± SD, median (range), or number (%), as appropriate. The biological unit was an individual mouse for animal studies and an independently performed experiment for cell studies. Statistical analyses were conducted using GraphPad Prism version 10. Two-group comparisons used unpaired two-tailed Student *t* tests or Mann-Whitney *U* tests as appropriate. Comparisons involving three or more groups used one-way ANOVA followed by Tukey-Kramer multiple-comparison testing; Dunnett testing was used for comparisons with a single control where indicated. Pearson correlation was used for the relationship between CD34-positive vascular profiles and FEV0.1/FVC. RNA-sequencing *P* values were adjusted using the Benjamini-Hochberg method. Exact group sizes and statistical tests are reported in the figure legends. *P* <0.05 was considered statistically significant.

## RESULTS

### HFD aggravates obstructive lung abnormalities in βENaC-Tg mice

We first asked whether the pulmonary response to HFD was influenced by a pre-existing muco-obstructive lung state. Male wild-type (WT) and βENaC-transgenic (βENaC-Tg) mice were maintained on a control diet (con) or HFD for 10–11 weeks beginning at 5–6 weeks of age (Fig. 1A). HFD increased body weight, fasting blood glucose, and total fat-pad mass in both genotypes (Fig. 1B–D). Longitudinal measurements of body weight and blood glucose, together with the weights of individual adipose depots, showed broadly similar measured systemic metabolic responses to HFD in WT and βENaC-Tg mice (Additional file 3: Fig. S1).

**Figure 1.**
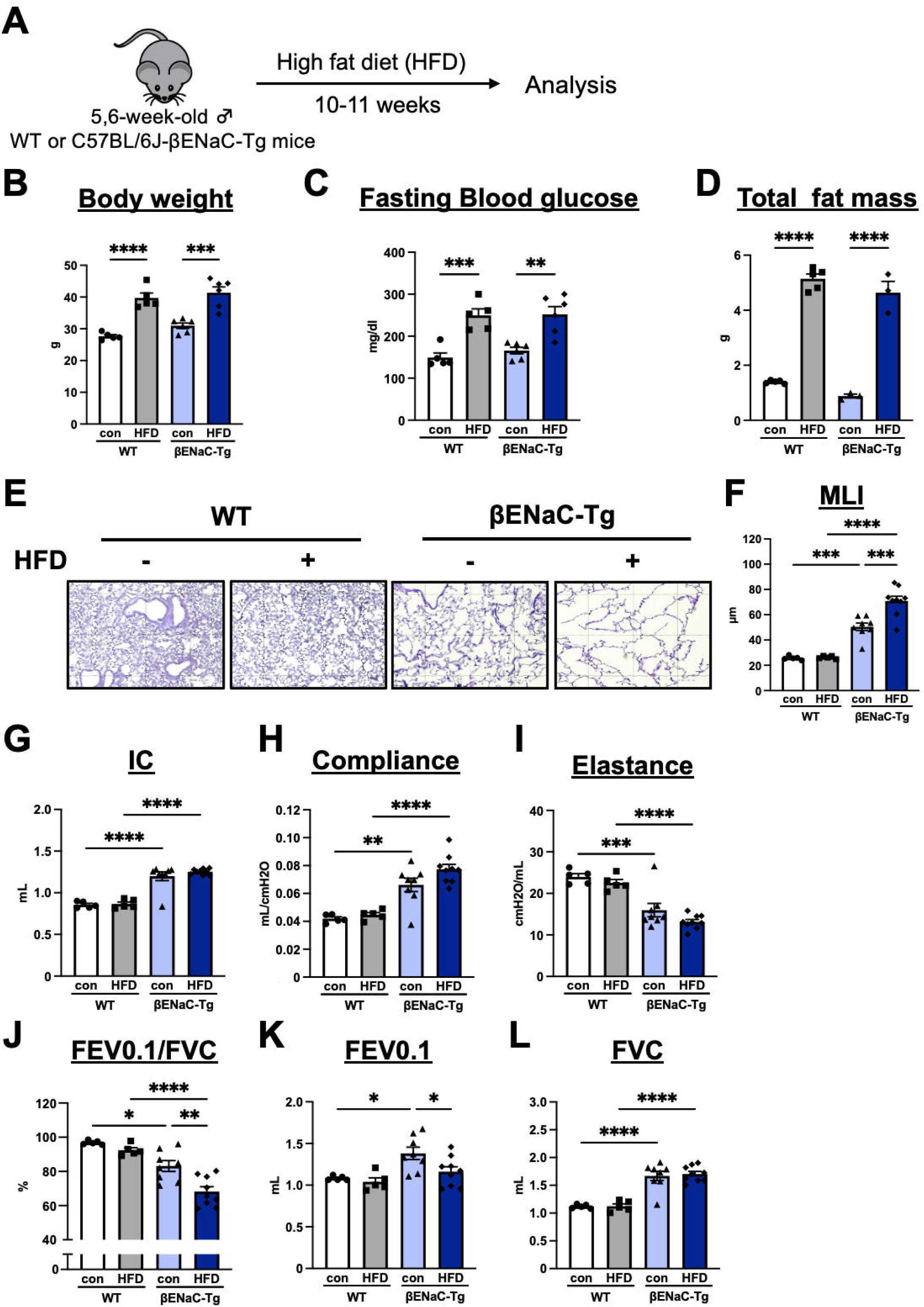
HFD aggravates obstructive lung abnormalities in βENaC-Tg mice. (A) Experimental design for male WT and βENaC-Tg mice fed control diet (CD) or high-fat diet (HFD) for 10-11 weeks beginning at 5-6 weeks of age. (B-D) Endpoint body weight, fasting blood glucose, and total fat-pad mass. (E) Representative H&E-stained lung sections. Scale bar, 100 μm. (F) Mean linear intercept (MLI). (G-I) Inspiratory capacity (IC), static compliance, and elastance measured using flexiVent. (J-L) FEV0.1/FVC, FEV0.1, and FVC. Data are mean ± SEM; each dot represents one mouse. Group sizes ranged from n = 5-9 mice per group according to endpoint. Findings were reproduced in an independent dietary cohort. Statistical significance was assessed by one-way ANOVA followed by Tukey-Kramer multiple-comparison testing. \**P* <0.05; \*\**P* <0.01; \*\*\**P* <0.001; \*\*\*\**P* <0.0001.

Under control diet conditions, βENaC-Tg mice already exhibited distal-airspace enlargement compared with WT mice. HFD further increased the mean linear intercept (MLI) in βENaC-Tg lungs (Fig. 1E,F). Additional morphometric indices, including alveolar area, perimeter, mean axis length, and Feret diameter, supported further airspace enlargement in HFD-fed βENaC-Tg mice (Additional file 3: Fig. S2).

Pulmonary-function analysis showed that βENaC-Tg mice had increased inspiratory capacity and static compliance and reduced elastance compared with WT mice, consistent with their baseline hyperinflation-like phenotype (Fig. 1G–I). These mechanical abnormalities were primarily associated with the βENaC-Tg disease background and were not markedly altered further by HFD. In contrast, HFD reduced FEV0.1/FVC in βENaC-Tg mice (Fig. 1J). Absolute FEV0.1 and FVC remained elevated in the setting of increased lung volumes and were therefore not interpreted as evidence of improved ventilatory function (Fig. 1K,L). Together, these findings show that HFD aggravates structural and expiratory abnormalities in the βENaC-Tg muco-obstructive lung disease model.

### HFD remodels lipid-metabolic, Akt-survival, and vascular programs in diseased lungs

To identify programs associated with the HFD response, we performed whole-lung RNA sequencing in WT and βENaC-Tg mice maintained on control diet or HFD. Under control-diet conditions, βENaC-Tg lungs already exhibited a distinct transcriptional state characterized by enhanced epithelial/secretory and innate immune programs, along with reduced stress-metabolic, matrix, and vascular signatures (Additional file 3: Fig. S3).

HFD induced broad transcriptional remodeling in both genotypes. In WT lungs, HFD reduced vascular-associated transcripts, including *Esm1*, *Gja5*, *Npr3*, and *Sox18*, and increased stress- and metabolic-response genes, including *Areg*, *Gdf15*, *Spp1*, *Cxcl2*, *Il17c*, *Hilpda*, *Fabp1*, *Apoc2*, *Hmgcs2*, *Mt1*, and *Mt2* (Additional file 3: Fig. S4). In βENaC-Tg lungs, HFD reduced genes linked to myeloid and innate immune programs, including *Retnlg*, *Mmp9*, *Cxcr2*, *Il36g*, *Trem3*, *Lst1*, *Siglece*, and *Cd209a*, while increasing genes associated with lipid handling, stress adaptation, and remodeling, including *Angptl4*, *Hmgcs2*, *Ucp2*, *Acot1*, *Sgk1*, *Fkbp5*, *Cebpb*, and *Lox* (Fig. 2A-C; Additional file 3: Fig. S5). Pathway analysis highlighted reduced PI3K-Akt-linked signaling as a prominent HFD-responsive node in βENaC-Tg lungs (Fig. 2D). These data indicate that the diseased lung enters metabolic challenge with a pre-remodeled transcriptional state and responds through coordinated survival, lipid-handling, immune, and vascular programs.

**Figure 2.**
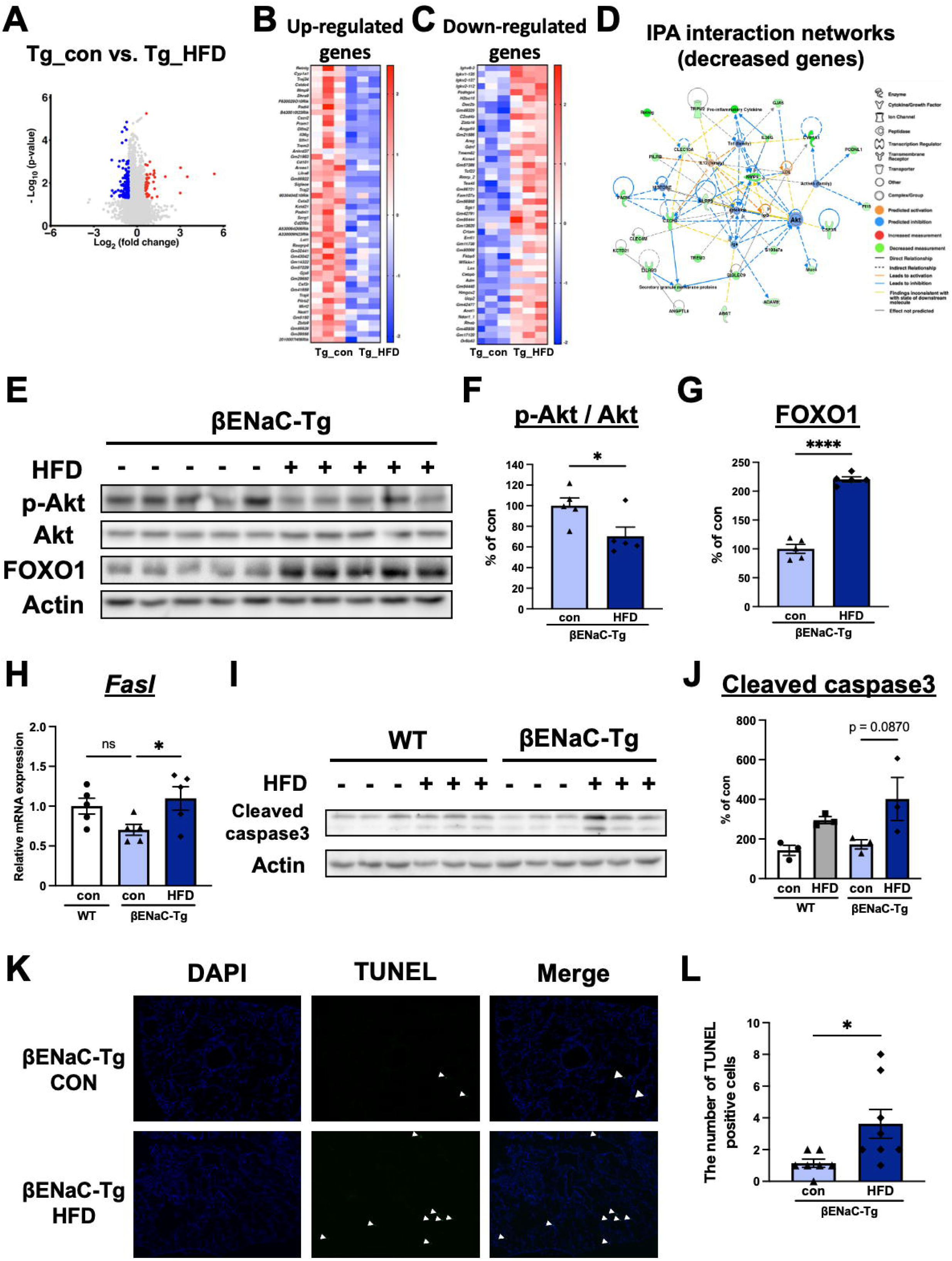
HFD remodels lung transcription and attenuates Akt survival signaling. (A) Volcano plot of differentially expressed genes (DEGs) in βENaC-Tg lungs after HFD versus CD. (B,C) Heatmaps of representative upregulated and downregulated genes; values are row z scores. (D) Ingenuity Pathway Analysis network generated from downregulated DEGs. (E) Immunoblots of phospho-Akt (p-Akt), total Akt, FOXO1, and actin. (F,G) Quantification of p-Akt/Akt and FOXO1 normalized to actin. (H) *Fasl* mRNA normalized to *18S rRNA*. (I,J) Cleaved caspase-3 immunoblot and quantification normalized to actin; the upward shift did not reach statistical significance (*P* = 0.0870). (K,L) Representative TUNEL staining and quantification of TUNEL-positive cells in epithelial regions. For RNA sequencing, n = 3 mice per group; DEGs were defined by absolute fold change >1.5 and FDR <0.05. Other group sizes ranged from n = 3-8 mice per group. Data are mean ± SEM; each dot represents one mouse. Two-group comparisons used unpaired two-tailed Student *t* tests; comparisons involving three or more groups used one-way ANOVA with Tukey-Kramer testing. ns, not significant; \**P* <0.05; \*\*\*\**P* <0.0001.

### HFD attenuates Akt signaling and enhances apoptosis-related responses in epithelial regions

We next examined whether the transcriptomic Akt signature was reflected at the protein and tissue levels. HFD reduced the p-Akt/Akt ratio and increased FOXO1 abundance in βENaC-Tg lungs (Fig. 2E-G). *Fasl* expression was increased, whereas cleaved caspase-3 abundance showed a nonsignificant upward trend (Fig. 2H–J). HFD also increased the number of TUNEL-positive nuclei within epithelial regions (Fig. 2K,L).

Together, these findings identify attenuation of Akt survival signaling, FOXO1 upregulation, and enhanced apoptosis-related activity in epithelial regions as components of the pulmonary response to HFD in βENaC-Tg mice.

### Short-term insulin-deficient hyperglycemia does not reproduce HFD-associated pulmonary injury

Because HFD produces combined lipid, glycemic, hormonal, and inflammatory changes, we examined whether short-term insulin-deficient hyperglycemia alone reproduced the pulmonary phenotype. Streptozotocin (STZ) administration produced sustained hyperglycemia and reduced serum insulin concentrations in βENaC-Tg mice (Fig. 3A,B). However, during the 3-week observation period, STZ did not increase MLI or produce apparent structural deterioration of the lung (Fig. 3C,D).

**Figure 3.**
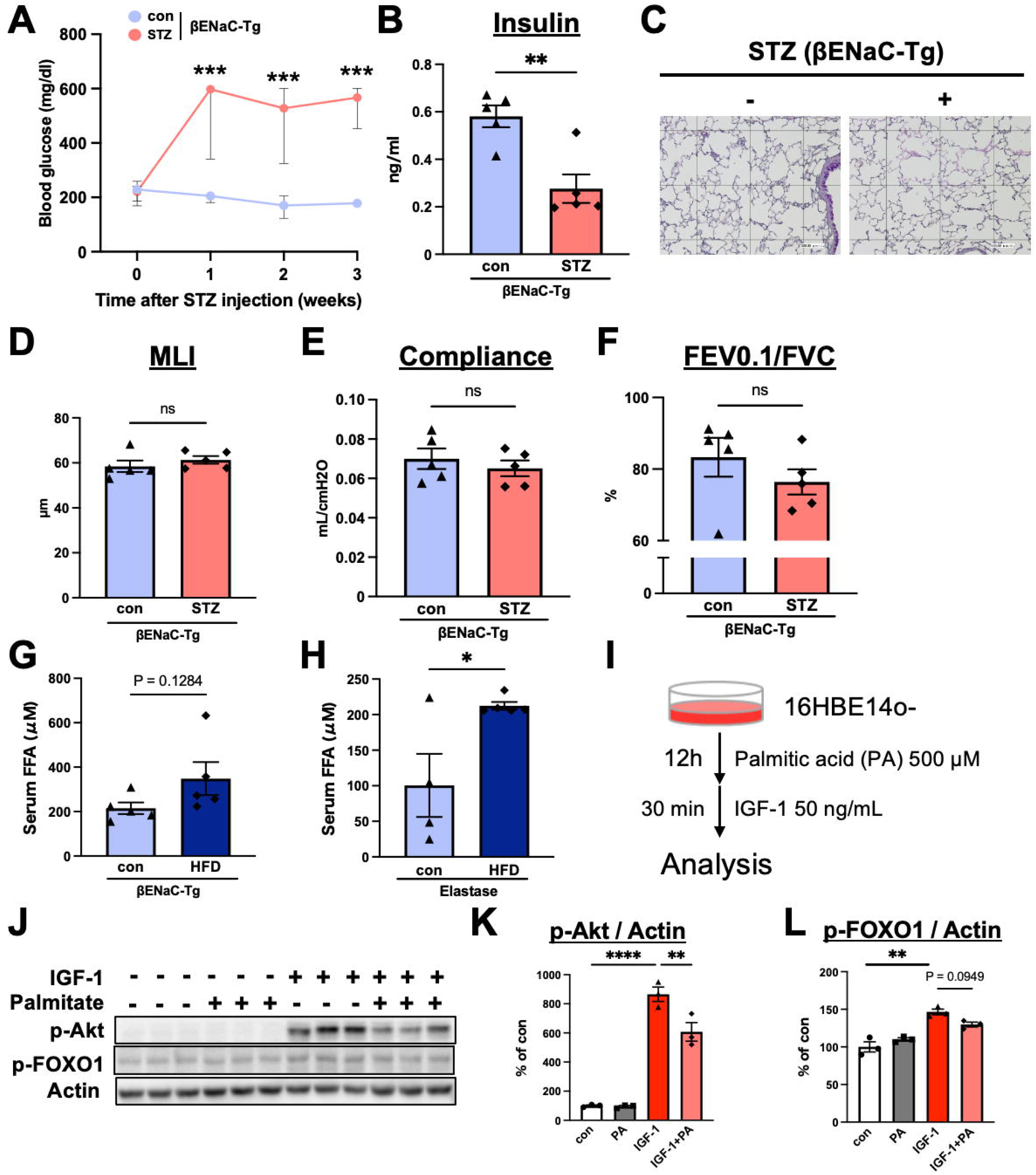
Lipid excess attenuates epithelial IGF-1-Akt responsiveness beyond hyperglycemia. (A) Blood-glucose time course after vehicle (con) or streptozotocin (STZ) administration in βENaC-Tg mice. (B) Endpoint serum insulin. (C) Representative PAS/Alcian blue-stained lung sections. (D) MLI. (E,F) Static compliance and FEV0.1/FVC. (G,H) Serum free-fatty-acid concentrations in βENaC-Tg mice and elastase-treated mice maintained on control diet (con) or HFD. (I) Palmitate/IGF-1 experimental design. Human bronchial epithelial 16HBE14o-cells were exposed for 12 h to palmitate-free control medium or palmitate-containing medium (500 μM) and then stimulated with IGF-1 (50 ng/mL) for 30 min. (J-L) Immunoblots and quantification of p-Akt and p-FOXO1 normalized to actin. *In vivo* group sizes ranged from n = 4-5 mice per group; cell experiments used n = 3 independent experiments per condition. Data are mean ± SEM. Statistical significance was assessed using unpaired two-tailed Student t tests or one-way ANOVA with Tukey-Kramer testing as appropriate. ns, not significant; \**P* <0.05; \*\**P* <0.01; \*\*\**P* <0.001; \*\*\*\**P* <0.0001.

Static compliance and FEV0.1/FVC were also unchanged after STZ treatment, and additional pulmonary-function indices showed no significant differences between the groups (Fig. 3E,F; Additional file 3: Fig. S6). Thus, short-term insulin-deficient hyperglycemia alone was insufficient to reproduce the structural and physiological changes observed after HFD exposure.

### Palmitate attenuates epithelial IGF-1-Akt responsiveness

Serum free-fatty-acid concentrations tended to be higher in HFD-fed βENaC-Tg mice and were significantly increased in HFD-fed elastase-treated mice (Fig. 3G,H). We therefore examined whether palmitate, as a representative saturated fatty acid, directly altered epithelial growth-factor signaling.

Human bronchial epithelial 16HBE14o-cells were exposed to palmitate-free control medium (CON) or palmitate-containing medium (PA; 500 μM sodium palmitate) for 12 h and subsequently stimulated with IGF-1 for 30 min (Fig. 3I). IGF-1 robustly increased Akt phosphorylation, whereas PA pretreatment markedly attenuated this response (Fig. 3J,K). IGF-1-induced FOXO1 phosphorylation showed a parallel but nonsignificant downward trend after PA pretreatment (*P* = 0.0949; Fig. 3J,L). Akt-pathway antibody-array profiling was consistent with broader attenuation of IGF-1-responsive phosphorylation signals involving Akt, PRAS40, GSK3α/β, p70 S6 kinase, and BAD after PA exposure (Additional file 3: Fig. S7). Together, these findings connect saturated-fatty-acid exposure to reduced epithelial responsiveness to IGF-1-Akt survival signaling.

### IGF-1R inhibition partially reproduces HFD-associated epithelial responses

To examine the functional contribution of IGF-1 signaling i*n vivo*, βENaC-Tg mice were treated with the IGF-1 receptor inhibitor picropodophyllin (PPP). PPP reduced pulmonary Akt phosphorylation, increased *Fasl* expression, and increased TUNEL-positive nuclei within epithelial regions (Fig. 4A–E). These changes reproduced key signaling and cellular features observed after HFD exposure. PPP produced smaller changes in MLI and pulmonary-function parameters than those observed with HFD, and these organ-level differences did not reach statistical significance (Fig. 4F–M). In elastase-treated mice, PPP similarly increased TUNEL positivity without significantly increasing airspace enlargement (Additional file 3: Fig. S8). These findings position impaired IGF-1-Akt signaling as one mechanistic component of a broader HFD-associated pulmonary injury response.

**Figure 4.**
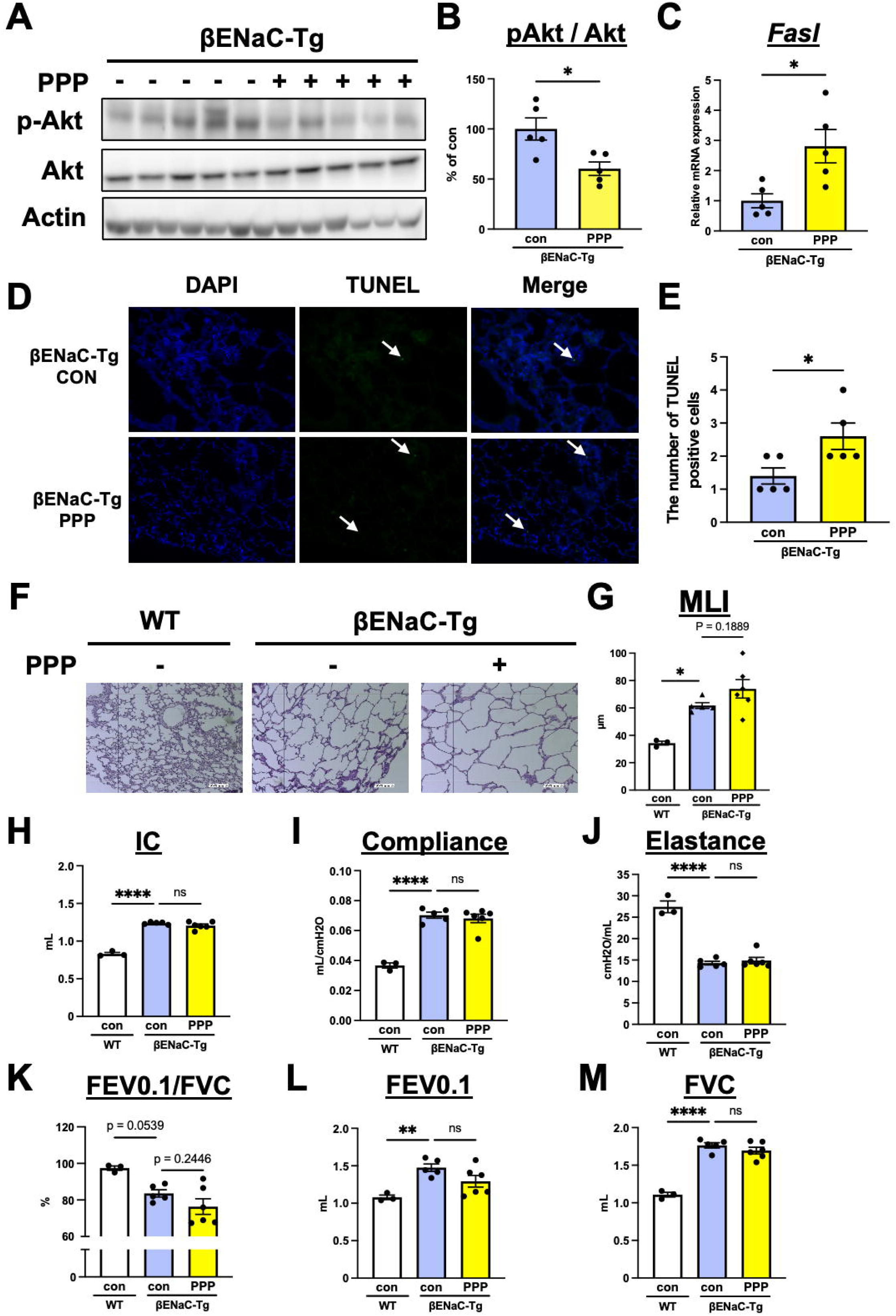
IGF-1R inhibition partially reproduces HFD-associated epithelial injury. (A) Immunoblots of p-Akt, total Akt, and actin after vehicle or picropodophyllin (PPP) treatment of βENaC-Tg mice. (B) Quantification of p-Akt/Akt. (C) *Fasl* mRNA normalized to *18S rRNA*. (D,E) Representative TUNEL staining and quantification in epithelial regions. (F,G) Representative lung histology and MLI. (H-M) Inspiratory capacity (IC), static compliance, elastance, FEV0.1/FVC, FEV0.1, and FVC. PPP was administered intraperitoneally at 60 μg/day for 14 days. Data are mean ± SEM; each dot represents one mouse. Group sizes ranged from n = 3-6 mice per group according to endpoint. Statistical significance was assessed using unpaired two-tailed Student *t* tests or one-way ANOVA with Tukey-Kramer testing as appropriate. ns, not significant; \**P* <0.05; \*\**P* <0.01; \*\*\*\**P* <0.0001.

### HFD preconditioning increases susceptibility to elastase-induced emphysema-like injury

We next asked whether an HFD-induced metabolic state also increased susceptibility to a distinct parenchymal insult. Mice were maintained on control diet or HFD for 10 weeks before intratracheal elastase administration (Fig. 5A). HFD increased body weight and fasting blood glucose and impaired glucose tolerance in elastase-treated mice (Fig. 5B-D).

**Figure 5.**
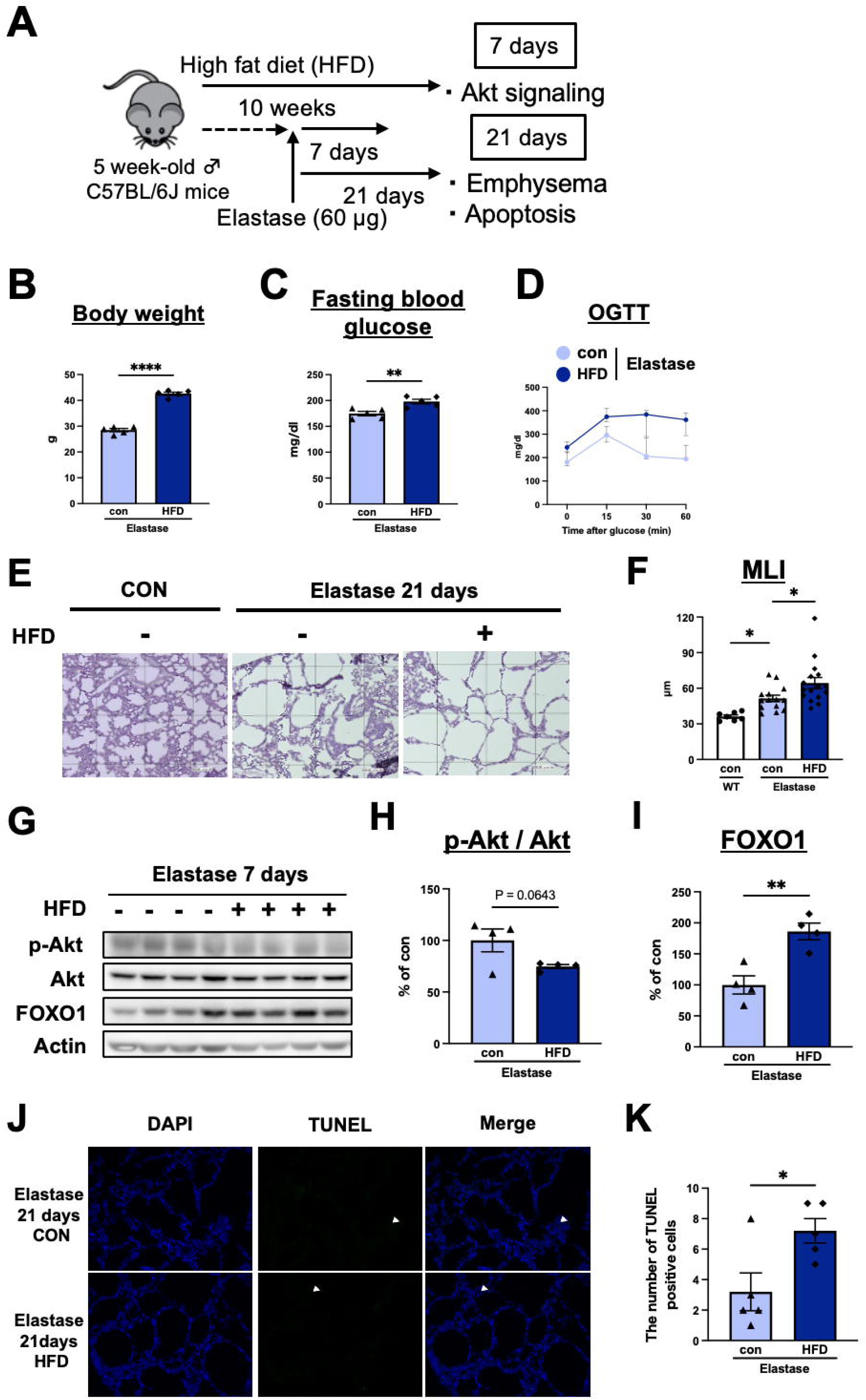
HFD preconditioning aggravates elastase-induced emphysema-like injury. (A) Experimental timeline. Male C57BL/6J mice were maintained on control diet (con) or HFD for 10 weeks before intratracheal elastase administration (60 μg). Akt-FOXO1 signaling was assessed 7 days after elastase; histology and TUNEL were assessed 21 days after elastase. (B-D) Body weight, fasting blood glucose, and oral glucose-tolerance responses. (E,F) Representative lung histology and MLI at day 21. (G-I) Immunoblots and quantification of p-Akt/Akt and FOXO1 normalized to actin at day 7; the reduction in p-Akt/Akt approached significance (*P* = 0.0643). (J,K) Representative TUNEL staining and quantification at day 21. Data are mean ± SEM; each dot represents one mouse. Group sizes ranged from n = 4-16 mice per group according to endpoint. Statistical significance was assessed using unpaired two-tailed Student *t* tests or one-way ANOVA with Tukey-Kramer testing as appropriate. \**P* <0.05; \*\**P* <0.01; \*\*\*\**P* <0.0001.

At 21 days after elastase, HFD-preconditioned mice displayed greater airspace enlargement and increased MLI (Fig. 5E,F). At 7 days after elastase, HFD was accompanied by a nonsignificant downward trend in the p-Akt/Akt ratio (*P* = 0.0643) and a significant increase in FOXO1 abundance. (Fig. 5G-I). At 21 days, TUNEL-positive cells were also increased (Fig. 5J,K). Thus, prior HFD exposure increased susceptibility to subsequent elastase-induced emphysema-like injury and was accompanied by Akt-FOXO1-associated survival changes similar in direction to those observed in the βENaC-Tg disease model.

### Lipid-stressed airway epithelium communicates injury signals to endothelial cells

The incomplete organ-level effect of IGF-1R inhibition, together with vascular signatures in the RNA-sequencing data, prompted us to examine the pulmonary endothelial compartment. CD34-positive vascular profiles were reduced in HFD-fed βENaC-Tg lungs (Fig. 6A,B), and vascular abundance positively correlated with FEV0.1/FVC across βENaC-Tg mice (Fig. 6C).

**Figure 6.**
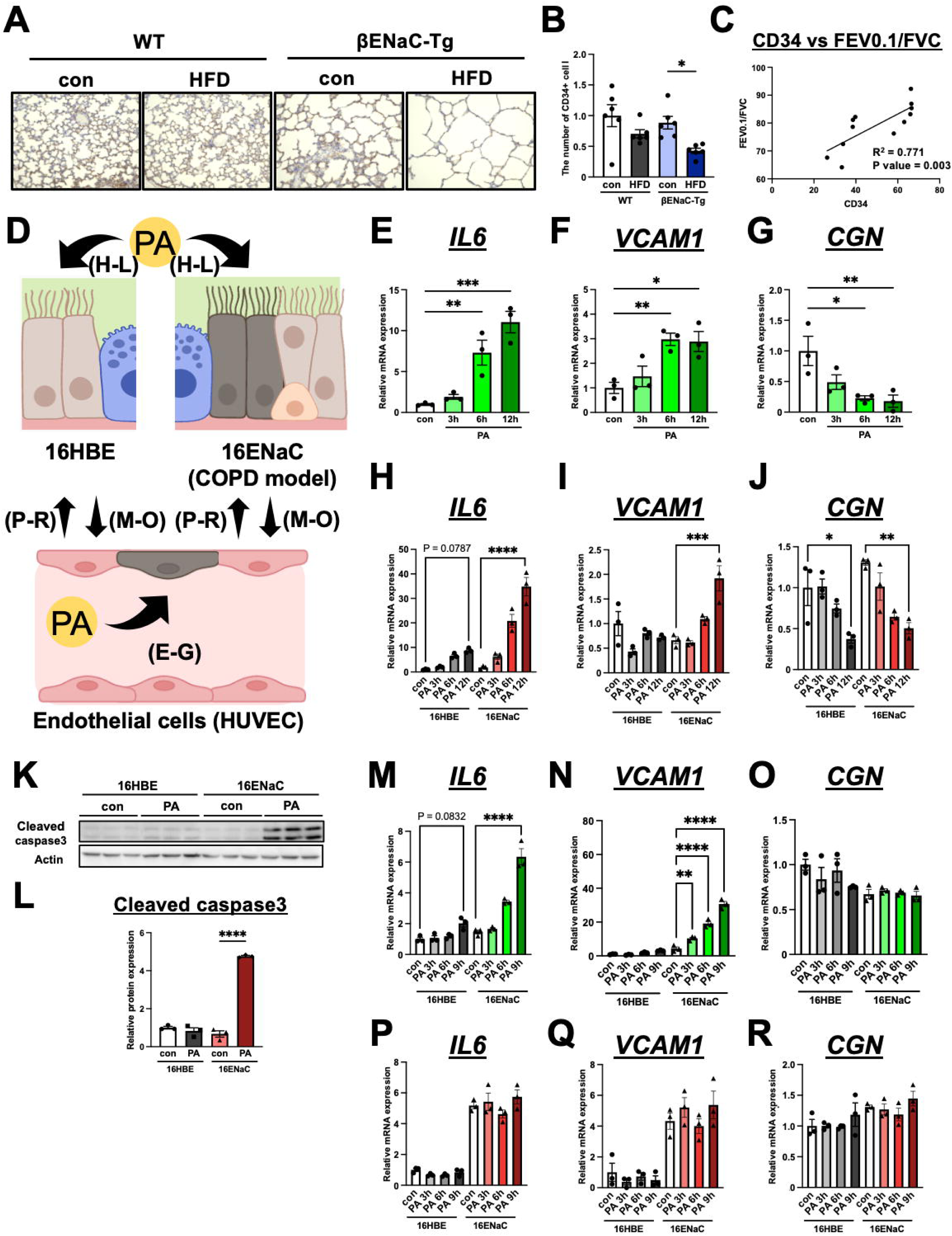
Lipid-stressed epithelium promotes endothelial stress responses. (A,B) Representative CD34 immunohistochemistry and quantification of CD34-positive vascular profiles in WT and βENaC-Tg mice maintained on CD or HFD. (C) Relationship between CD34-positive vascular profiles and FEV0.1/FVC in βENaC-Tg mice assessed by two-tailed Pearson correlation. (D) Experimental scheme for direct palmitate exposure and bidirectional conditioned-medium transfer using parental 16HBE14o-cells, ENaC-hyperactive 16HBE14o-β/γENaC cells, and HUVECs. (E-G) Direct effects of palmitate on endothelial *IL6*, *VCAM1*, and *CGN* expression. (H-L) Palmitate responses in parental and ENaC-hyperactive 16HBE14o-airway epithelial cells, assessed by *IL6*, *VCAM1*, *CGN*, and cleaved caspase-3. (M-O) HUVEC responses to conditioned media from palmitate-exposed epithelial cells. (P-R) Epithelial responses to conditioned media from palmitate-exposed HUVECs. Donor cells were exposed to palmitate-containing medium (500 μM) or palmitate-free control medium for 3 h; conditioned media were collected 3, 6, and 9 h after medium replacement and applied to recipient cells for 30 min. For (B), n = 6 mice per group; the correlation in (C) included 12 βENaC-Tg mice. Cell experiments used n = 3 independent experiments per condition. Data are mean ± SEM. One-way ANOVA with Tukey-Kramer testing was used for group comparisons. \**P* <0.05; \*\**P* <0.01; \*\*\**P* <0.001; \*\*\*\**P* <0.0001.

Palmitate (PA) directly increased *IL6* and *VCAM1* and reduced the junction-associated gene *CGN* in HUVECs (Fig. 6D-G). ENaC-hyperactive epithelial cells also mounted a pronounced response to palmitate, with induction of *IL6* and *VCAM1*, lower *CGN* expression, and increased cleaved caspase-3 (Fig. 6H-L).

To test directional communication, epithelial cells or HUVECs were transiently exposed to palmitate, the treatment medium was replaced, and conditioned media were transferred to the reciprocal cell type. Conditioned media from palmitate-exposed epithelial cells induced endothelial *IL6* and *VCAM1* expression, and *CGN* expression showed a modest downward trend, with the strongest responses observed using ENaC-hyperactive epithelial donors (Fig. 6M-O). In contrast, conditioned media from palmitate-exposed HUVECs produced limited responses in epithelial recipient cells (Fig. 6P-R). These data reveal a predominantly epithelial-to-endothelial signaling pattern in which an ENaC-hyperactive epithelial state amplifies the propagation of lipid stress.

### Hepatic steatosis is associated with lower percent-predicted FEV1 in men with airflow obstruction

We finally examined whether a metabolically adverse phenotype was associated with greater lung-function impairment in humans. The cohort included 474 Japanese men, comprising 185 never-smokers and 289 ever-smokers, with a mean age of 63.3 ± 12.4 years (Table 1). Hepatic steatosis was present in 22.8% of participants, and airflow obstruction was present in 16.0%.

**Table 1.** Clinical characteristics of human male participants.

|  | All<br>(n = 474) | Smoking status |  | P |
| --- | --- | --- | --- | --- |
|  |  | Never-smoker<br>(n = 185) | Ever-smoker<br>(n = 289) |  |
| Age (years) | 63.3 ± 12.4 | 64.9 ± 12.6 | 62.3 ± 12.2 | 0.026 |
| BMI (kg/m<sup>2</sup>) | 23.4 ± 2.7 | 23.3 ± 2.7 | 23.4 ± 2.8 | 0.797 |
| BMI ≥ 23 kg/m<sup>2</sup> (%) | 268 (56.5) | 101 (54.6) | 167 (57.8) | 0.278<sup>a</sup> |
| BMI ≥ 25 kg/m<sup>2</sup> (%) | 131 (27.6) | 51 (27.6) | 80 (27.7) | 0.999<sup>a</sup> |
| FBG (mg/dL) | 98 (75 – 244) | 97 (77 – 206) | 99 (75 – 244) | 0.325<sup>b</sup> |
| HDL-C (mg/dL) | 62.6 ± 15.3 | 63.5 ± 14.2 | 62.0 ± 16.1 | 0.290 |
| LDL-C (mg/dL) | 119.0 ± 26.5 | 117.9 ± 25.4 | 119.6 ± 27.2 | 0.503 |
| TGs (mg/dL) | 95 (26 – 520) | 90 (31 – 302) | 98 (26 – 520) | 0.012<sup>b</sup> |
| AST (IU/L) | 25.5 ± 10.6 | 25.1 ± 8.3 | 25.7 ± 11.9 | 0.550 |
| ALT (IU/L) | 25.0 ± 15.0 | 24.4 ± 15.6 | 25.4 ± 14.5 | 0.476 |
| GGT (IU/L) | 29 (6 – 657) | 27 (10 – 185) | 30 (6 – 657) | 0.163<sup>b</sup> |
| FVC (ml) | 3707 ± 693 | 3651 ± 662 | 3742 ± 711 | 0.161 |
| FEV1 (ml) | 2796 ± 620 | 2795 ± 606 | 2796 ± 629 | 0.984 |
| FEV1/FVC (%) | 75.3 ± 7.8 | 76.4 ± 6.7 | 74.6 ± 8.4 | 0.015 |
| Percent predicted FEV1 (%) | 92.7 ± 14.7 | 95.3 ± 13.9 | 91.0 ± 15.0 | 0.002 |
| Airflow obstruction (%) | 76 (16.0) | 19 (10.3) | 57 (19.7) | 0.007<sup>a</sup> |
| Fatty liver (%) | 108 (22.8) | 38 (20.5) | 70 (24.2) | 0.371<sup>a</sup> |
| Diabetes (%) | 87 (18.4) | 29 (15.7) | 58 (20.1) | 0.274<sup>a</sup> |
| Hypertension (%) | 201 (42.4) | 88 (47.6) | 113 (39.1) | 0.071<sup>a</sup> |
| Smoking period (years) | — | — | 25.8 ± 13.1 | — |
| Brinkman index | — | — | 601 ± 429 | — |
| Alcohol intake (g/day) | 10 (0 – 90) | 5 (0 – 60) | 14.3 (0 – 90) | 0.004<sup>b</sup> |

Values are mean ± standard deviation, median (range), or number of subjects (%).
<sup>a</sup> Fisher's exact test. <sup>b</sup> Mann-Whitney U test (otherwise, Student's *t*-test was used).
BMI, body mass index; FBG fasting blood glucose; HDL-C, high-density lipoprotein
cholesterol; LDL-C, low-density lipoprotein cholesterol; TGs, triglycerides; AST,
aspartate aminotransferase; ALT, alanine aminotransferase; GGT, $\gamma$ -glutamyltransferase.

Among participants with airflow obstruction, those with hepatic steatosis had lower percent-predicted FEV1 than those without hepatic steatosis (Fig. 7A). The same pattern was observed when the analysis was restricted to ever-smokers (Fig. 7B). BMI-based sensitivity analyses using thresholds of 25 and 27 kg/m² were less consistent and are presented separately in Additional file 3: Fig. S9. These findings provide a clinical context in which hepatic steatosis accompanies greater functional impairment among men with airflow obstruction.

**Figure 7.**
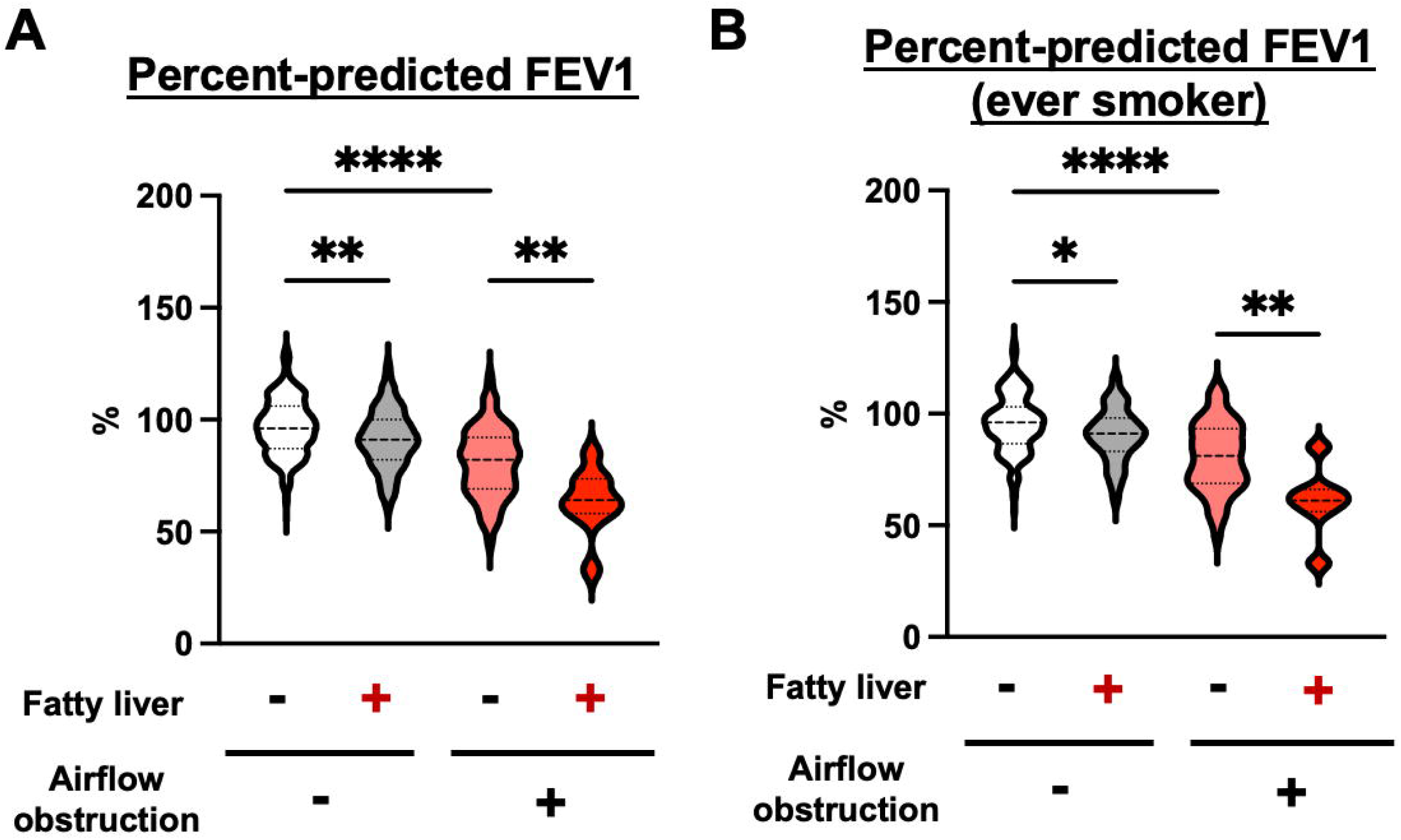
Hepatic steatosis is associated with lower percent-predicted FEV1 in men with airflow obstruction. (A) Percent-predicted FEV1 stratified according to hepatic steatosis and airflow-obstruction status in the entire cohort. From left to right, the groups comprised participants without hepatic steatosis or airflow obstruction (n = 299), with hepatic steatosis but without airflow obstruction (n = 99), without hepatic steatosis but with airflow obstruction (n = 67), and with both hepatic steatosis and airflow obstruction (n = 9). (B) The corresponding analysis restricted to ever-smokers. From left to right, group sizes were n = 169, 63, 50, and 7, respectively. Airflow obstruction was defined as a pre-bronchodilator FEV1/FVC ratio <0.70. Hepatic steatosis was assessed by ultrasonography. Violin plots show the distribution of individual values, and horizontal bars indicate medians. Statistical significance was assessed using one-way ANOVA followed by Tukey–Kramer multiple-comparison testing. \**P* <0.05; \*\**P* <0.01; \*\*\*\**P* <0.0001.

## Discussion

This study identifies the obstructive lung as a metabolically vulnerable organ. HFD increased body weight, glycemia, and adiposity in both WT and βENaC-Tg mice, but amplified distal-airspace enlargement and expiratory dysfunction within the βENaC-Tg disease background. Whole-lung transcriptomics and tissue validation linked this response to coordinated remodeling of lipid-handling, PI3K-Akt survival, and vascular programs. Palmitate reduced epithelial IGF-1-Akt responsiveness, IGF-1R inhibition reproduced the signaling and apoptosis-related component of the response, and lipid-exposed ENaC-hyperactive epithelium transmitted stress signals to endothelial cells. HFD preconditioning also increased susceptibility to elastase-induced emphysema-like injury. Together, these findings support a model in which systemic lipid stress interacts with pulmonary disease context to engage a multicompartment epithelial-vascular injury program.

Recent studies have established macrophage-centered mechanisms of lipid-associated COPD, including PLA2G7-arachidonic acid-NLRP3 signaling during severe diet-induced obesity [17], cholesterol-driven mitochondrial stress [18], and ceramide transfer from macrophages to endothelial cells [19]. Airway epithelial extracellular vesicles can also influence the vascular compartment in COPD [20]. Our findings extend this field in a complementary direction. Rather than asking whether severe obesity can initiate lung injury, we asked how an established muco-obstructive lung responds to systemic metabolic stress. The combination of a chronic βENaC-associated airway phenotype, a palmitate-sensitive epithelial IGF-1-Akt response, and epithelial-to-endothelial conditioned-medium activity defines a structural-cell dimension of pulmonary lipotoxicity.

The βENaC-Tg model was informative because mucus obstruction, epithelial stress, inflammation, and distal-airspace abnormalities were already present before HFD exposure [26–28]. HFD elicited similar measured systemic metabolic changes in WT and βENaC-Tg mice, yet the disease model developed additional airspace enlargement and a lower FEV0.1/FVC. This pattern supports the concept that tissue context, not systemic exposure alone, determines the pulmonary consequence of metabolic stress. The baseline transcriptomic state of βENaC-Tg lungs further suggests that epithelial, immune, matrix, and vascular programs are reorganized before the dietary challenge, creating a permissive state for HFD-induced injury.

Akt emerged as a central survival node within this response. Akt coordinates epithelial homeostasis [34], mitochondrial survival [35], and protection against oxidant lung injury [36]. HFD reduced Akt phosphorylation and increased FOXO1, *Fasl*, and TUNEL positivity in epithelial regions. Palmitate then provided a direct link between lipid exposure and impaired growth-factor responsiveness by attenuating IGF-1-induced Akt activation and multiple downstream pathway nodes. Saturated membrane lipids can activate endoplasmic-reticulum stress [37], and palmitate can promote calcium depletion, mitochondrial oxidative stress, and apoptosis [38]. A palmitate-rich HFD also exacerbates experimental pulmonary fibrosis [39], supporting the broader capacity of saturated-fatty-acid stress to modify lung injury.

The PPP experiments place IGF-1-Akt signaling within, rather than above, the broader HFD response. IGF-1R inhibition reproduced Akt suppression, *Fasl* induction, and increased TUNEL positivity, but produced smaller organ-level effects than HFD. This pattern is consistent with a partial mechanistic contribution of epithelial survival failure alongside additional lipid-responsive pathways. It also fits the context-dependent biology of IGF-1 in the lung: IGF-1 can support pulmonary growth and vascular development [40], whereas IGF-1R blockade can be beneficial in selected injury settings [41]. The present data therefore identify impaired epithelial IGF-1-Akt responsiveness as a component of metabolic vulnerability and point to a broader network of structural and inflammatory responses.

The elastase experiment strengthens the general relevance of this concept. Here, HFD preceded rather than followed the emphysema-inducing insult. HFD preconditioning increased subsequent airspace enlargement and was accompanied by lower Akt activity, FOXO1 upregulation, and greater TUNEL positivity. Thus, the two models address complementary scenarios: metabolic stress superimposed on chronic muco-obstructive disease in βENaC-Tg mice, and metabolic priming before an acquired parenchymal injury in the elastase model. Their convergence suggests that lipid-associated survival failure can operate across anatomically distinct COPD-relevant contexts.

The vascular findings add a second structural-cell arm. Pulmonary endothelial integrity is essential for alveolar maintenance, and disruption of VEGF-receptor signaling can induce endothelial apoptosis and emphysema [42]. HFD reduced CD34-positive vascular profiles in βENaC-Tg lungs, while palmitate directly induced endothelial inflammatory and adhesion programs and reduced *CGN* expression. These responses are consistent with established effects of free fatty acids on endothelial signaling [43] and with ANGPTL4-mediated lipid-responsive regulation of endothelial junctions [44].

Conditioned-medium transfer further showed that the epithelium can propagate lipid stress to the endothelium. The strongest endothelial responses arose from palmitate-exposed ENaC-hyperactive epithelial cells, linking the magnitude of intercellular signaling to the pre-existing epithelial state. This finding complements macrophage-to-endothelial ceramide transfer [19] and epithelial-endothelial communication described in other lung-injury settings [45]. The resulting model is not a single-cell lipotoxic pathway, but a network in which epithelial survival signaling and vascular homeostasis are jointly destabilized by metabolic stress.

The human analysis provides a clinically relevant counterpart. Longitudinal studies have linked increasing adiposity and progression of fatty liver to lung-function decline [46,47], and metabolic dysfunction-associated steatotic liver disease is increasingly recognized in the respiratory comorbidity landscape [48,49]. In the present cohort, hepatic steatosis coincided with lower percent-predicted FEV1 among men with airflow obstruction, including ever-smokers. This cross-sectional association supports the translational premise that ectopic lipid accumulation may more effectively mark a metabolically vulnerable obstructive lung phenotype than conventional overweight alone.

Several limitations define the next stage of this work. The study used male mice, bulk RNA sequencing with three samples per group, and HFD as a complex systemic exposure; comprehensive lipidomics will be required to identify the responsible lipid species. The epithelial and endothelial culture systems do not capture the full diversity of differentiated airway and pulmonary vascular cells, and the epithelial-derived mediator remains to be identified. The clinical cohort was retrospective, male-only, and based on pre-bronchodilator airflow obstruction. These considerations motivate cell-specific lipid profiling, primary epithelial-pulmonary endothelial co-culture, spatial analysis, and targeted rescue experiments. Single-cell atlases provide a framework for resolving the participating compartments [50].

## Conclusions

Our findings position dietary lipid stress as a modifier of COPD-relevant lung injury and support the concept that the obstructive lung is metabolically vulnerable. HFD-associated injury converged on impaired epithelial IGF-1-Akt signaling, FOXO1-associated apoptosis-related activity, vascular alterations, and epithelial-endothelial communication, supporting a multicompartment model of pulmonary lipotoxicity. The accompanying association between hepatic steatosis and lower lung function in individuals with airflow obstruction provides clinical context and a rationale for defining lipid- and cell-specific drivers of metabolically vulnerable COPD.

## Supporting information

uncropped blots

## List of abbreviations

BMI: body mass index
BSA: bovine serum albumin
COPD: chronic obstructive pulmonary disease
FFA: free fatty acid
FEV0.1: forced expiratory volume in 0.1 s
FEV1: forced expiratory volume in 1 s
FVC: forced vital capacity
HFD: high-fat diet
HUVEC: human umbilical vein endothelial cell
IGF-1: insulin-like growth factor-1
IGF-1R: insulin-like growth factor-1 receptor
MASLD: metabolic dysfunction-associated steatotic liver disease
MLI: mean linear intercept
PA: palmitate
PI3K: phosphatidylinositol-3-kinase
PPP: picropodophyllin
ROS: reactive oxygen species
STZ: streptozotocin
TPM: transcripts per million
WT: wild type

## Declarations

## Ethics approval and consent to participate

All animal procedures were approved by the Animal Welfare Committee of Kumamoto University (approval no. A2024-052) and were conducted in accordance with institutional and national guidelines. The human study was conducted in accordance with the Declaration of Helsinki and was approved by the ethics committees of the Japanese Red Cross Kumamoto Health Care Center and the Faculty of Life Sciences, Kumamoto University (Genome no. 169). All participants provided written informed consent.

## Consent for publication

Not applicable.

## Availability of data and materials

The RNA-sequencing dataset generated during the current study is available in the NCBI Gene Expression Omnibus under accession GSE300375 [51]. Individual-level clinical screening data are not publicly available because of participant privacy and ethics restrictions but may be available from the corresponding author on reasonable request, subject to approval by the relevant ethics committees and an appropriate data-use agreement. Materials generated in this study are available from the corresponding author on reasonable request and completion of an appropriate material-transfer agreement.

## Authors’ contributions

C.O., H.N., T.K., and T.S. conceived and designed the study. C.O., H.N., T.K., K.U-S., and T.S. performed and analyzed animal and cell-culture experiments. C.O., K.O., J.S., Y.O., and T.S. performed and analyzed the human study. C.O., T.K., K.W., Y.M., and T.S. performed and analyzed RNA-sequencing experiments. C.O., H.N., T.K., K.K., and Y.F. performed and analyzed immunohistochemical studies. C.O., H.N., T.K., K.K., R.N., S.K., N.T., M.H., A.F., and M.U. contributed to mouse experiments. C.O., H.N., T.K., M.A.S., and T.S. drafted and revised the manuscript. H.K. and T.S. supervised the study. All authors read and approved the final manuscript.

## Acknowledgements

The authors thank the staff of the Center for Animal Resources and Development and the research-support facilities at Kumamoto University for technical assistance.

## Authors’ information

Not applicable.

## Additional files

**Additional file 1.** DOCX. Supplementary Table S1. Oligonucleotide sequences used for quantitative RT-PCR.

**Additional file 2.** PDF. Uncropped immunoblots corresponding to the main and supplementary figures.

**Additional file 3.** PDF. Supplementary Figures S1–S9 and their legends.

