## Supplementary material for "Dietary lipid stress aggravates obstructive lung injury involving epithelial IGF-1-Akt dysfunction and epithelial-endothelial crosstalk": Table. S1

**Table S1. Oligonucleotide sequences used for quantitative RT-PCR.**

| Primer | Sequence |
| --- | --- |
| m18S rRNA_QRT-FW | 5′–GTAACCCGTTGAACCCCATT–3′ |
| m18S rRNA_QRT-RV | 5′–CCATCCAATCGGTAGTAGCG–3′ |
| mFasl_QRT-FW | 5′–TCCGTGAGTTCACCAACCAAA–3′ |
| mFasl_QRT-RV | 5′–GGGGGTTCCCTGTTAAATGGG–3′ |
| hIL-6_QRT-FW | 5′–GCACTGGCAGAAAACAACCT–3′ |
| hIL-6_QRT-RV | 5′–CAGGGGTGGTTATTGCATCT–3′ |
| hVCAM1_QRT-FW | 5′–GATTCTGTGCCCACAGTAAGGC–3′ |
| hVCAM1_QRT-RV | 5′–TGGTCACAGAGCCACCTTCTTG–3′ |
| hCGN_QRT-FW | 5′–CAAGGAGGATCTTAGAGCCACC–3′ |
| hCGN_QRT-RV | 5′–TGGCGAGTATCTCCAGCACTAG–3′ |
| hGAPDH_QRT-FW | 5′–TCCACTGGCGTCTTCACC–3′ |
| hGAPDH_QRT-RV | 5′–GGCAGAGATGATGACCCTTTT–3′ |
