## Supplementary Figures S1-S9 for "Dietary lipid stress aggravates obstructive lung injury involving epithelial IGF-1-Akt dysfunction and epithelial-endothelial crosstalk"

Supplementary Fig. S1-S9

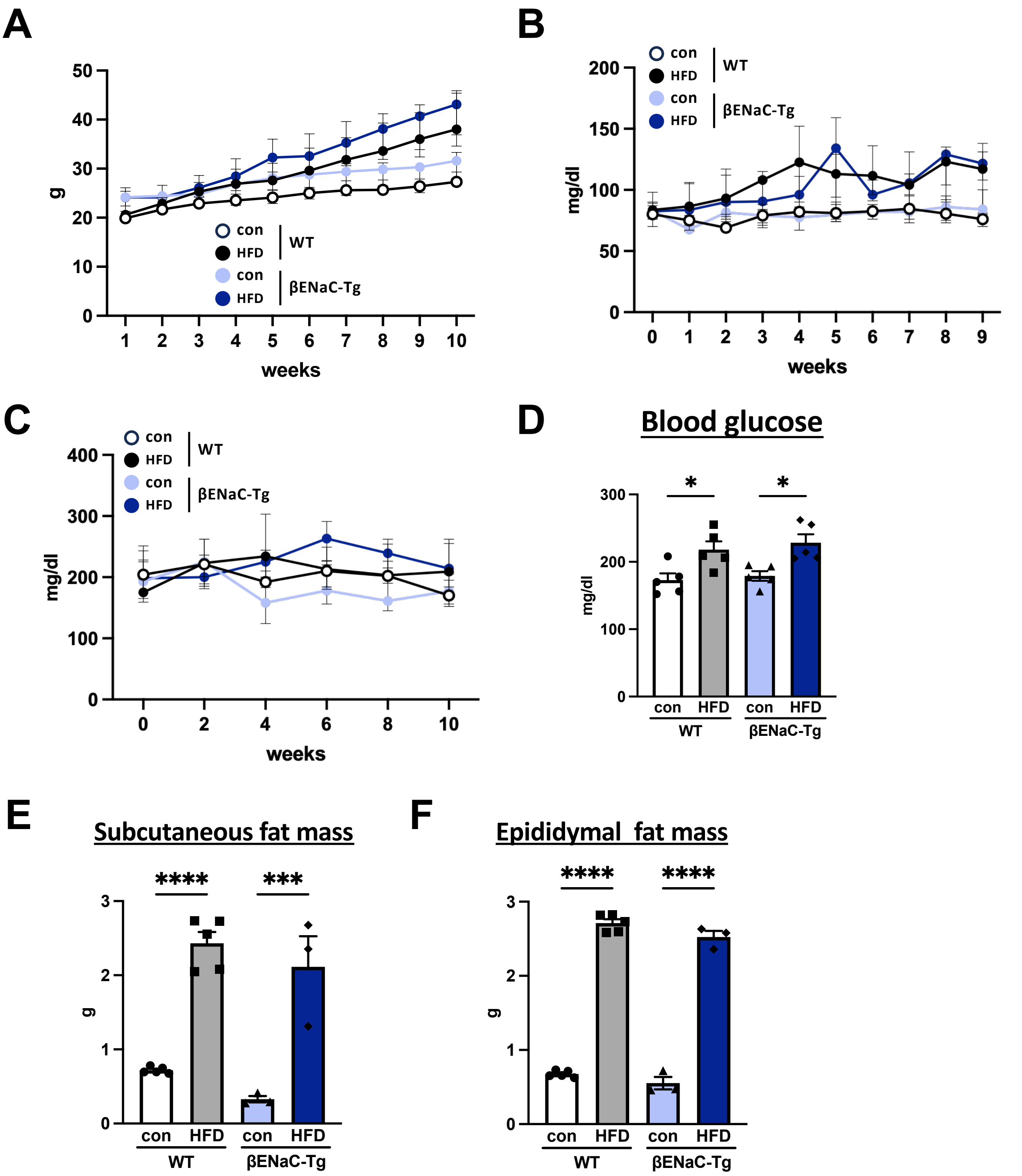


**Supplementary Figure S1. HFD induces similar systemic metabolic responses in WT and βENaC-Tg mice.** (A-C) Body-weight, fasting-glucose, and non-fasting-glucose time courses. (D) Endpoint non-fasting blood glucose. (E,F) Subcutaneous and epididymal fat-pad mass. Data are mean ± SEM; each dot represents one mouse. n = 5-8 mice per group. One-way ANOVA with Tukey multiple-comparison testing was used for (D-F). *P <0.05; **P <0.01; ***P <0.001; ****P <0.0001.

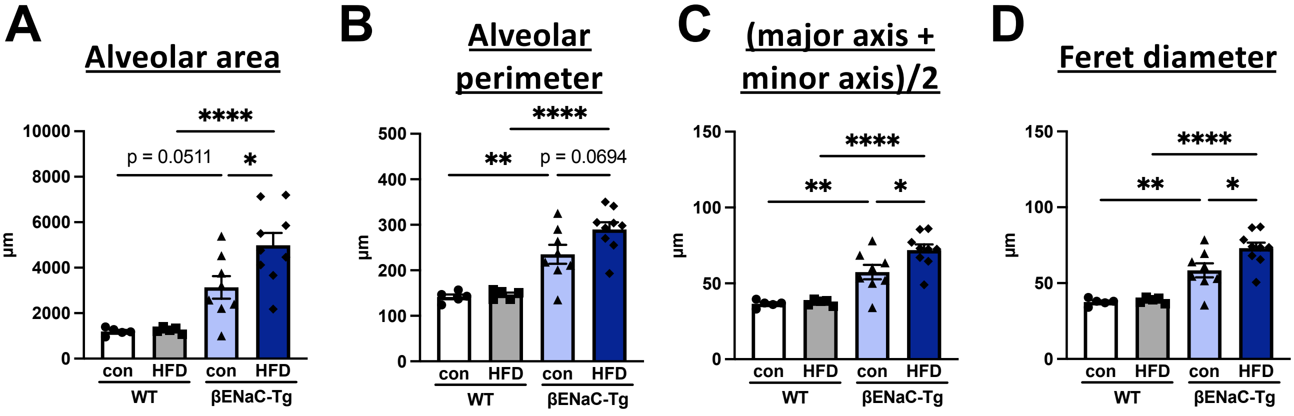


**Supplementary Figure S2. Additional morphometry confirms HFD-associated airspace enlargement in βENaC-Tg mice.** (A-D) Alveolar area, alveolar perimeter, mean of major and minor axes, and Feret diameter. Data are mean ± SEM; each dot represents one mouse. n = 5-8 mice per group. One-way ANOVA with Tukey multiple-comparison testing was used. *P <0.05; **P <0.01; ***P <0.001; ****P <0.0001.

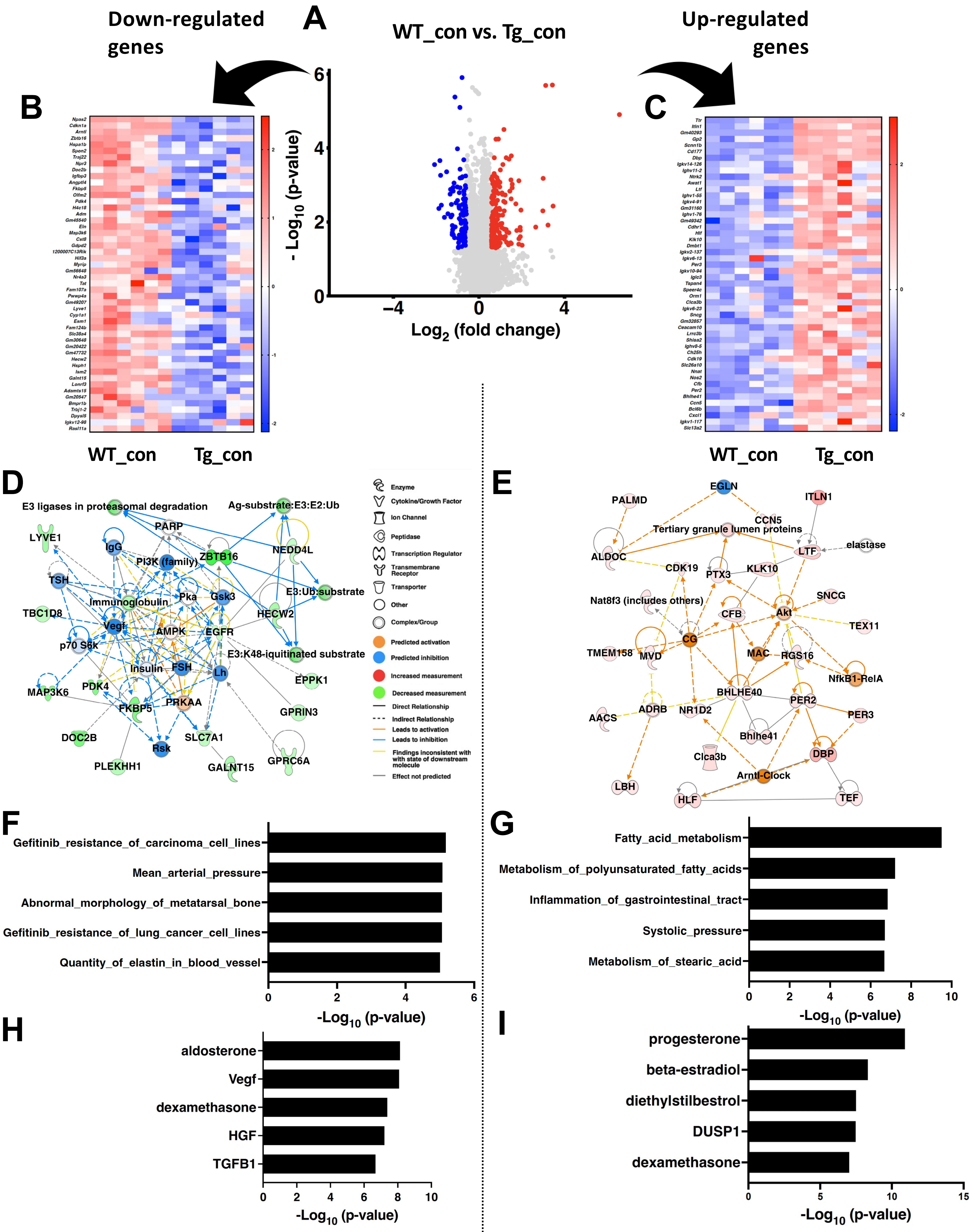


**Supplementary Figure S3. Baseline transcriptional remodeling in βENaC-Tg lungs.** (A) Volcano plot of DEGs comparing WT_con versus Tg_con. (B,C) Heatmaps of representative DEGs decreased (B) or increased (C) in Tg_con versus WT_con (row z-scores). (D,E) IPA interaction networks generated from DEGs decreased (D) or increased (E). (F,G) IPA diseases and functions enriched among decreased (F) or increased (G) DEGs (−log10 P values). (H,I) IPA upstream regulator analysis for decreased (H) or increased (I) DEGs. DEGs were defined by absolute fold change > 1.5 and FDR < 0.05 (n = 3 mice per group).

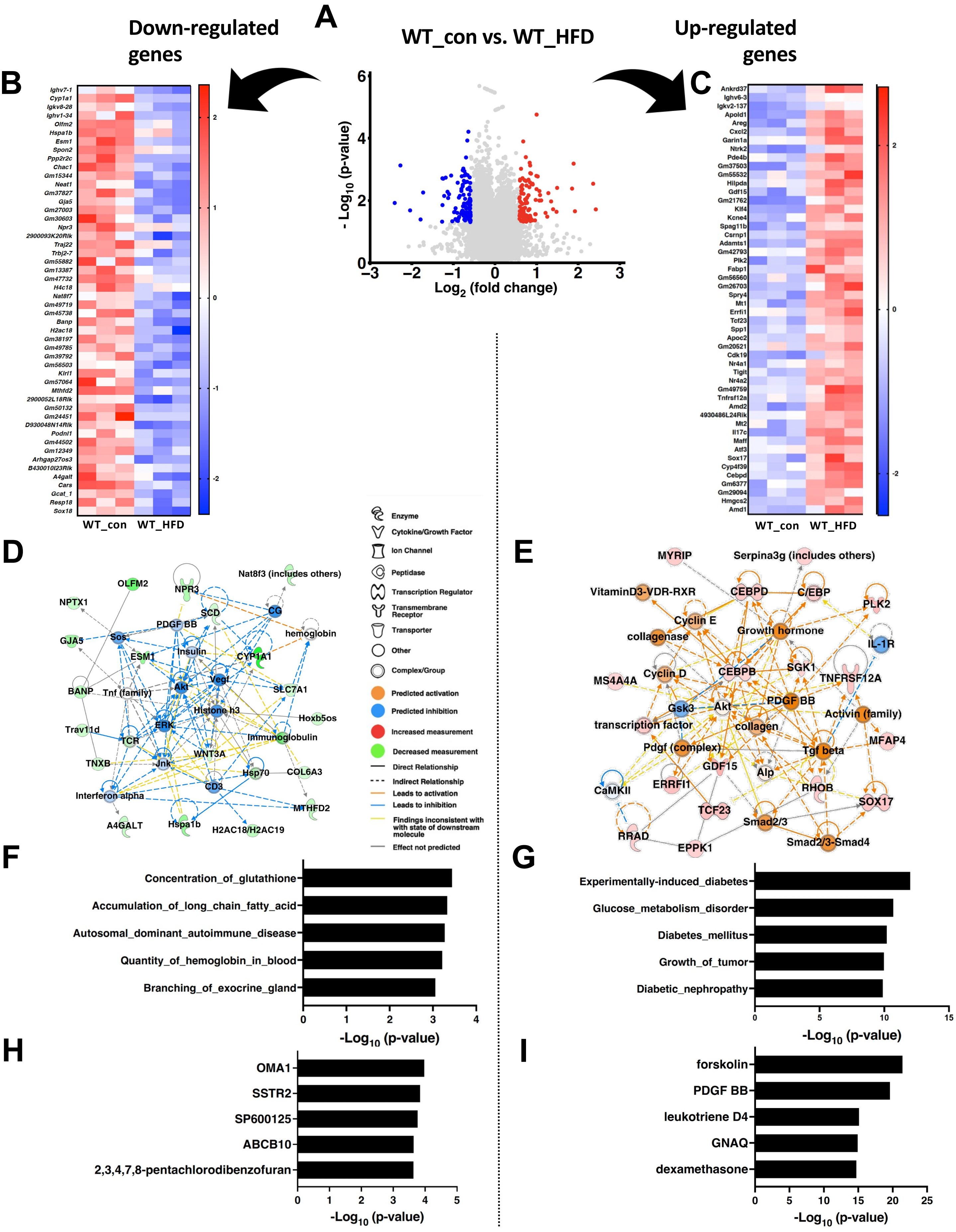


**Supplementary Figure S4. HFD induces transcriptional remodeling in WT lungs.** (A) Volcano plot of DEGs comparing WT_con versus WT_HFD. (B,C) Heatmaps of representative DEGs decreased (B) or increased (C) in WT_HFD versus WT_con (row z-scores). (D,E) IPA interaction networks generated from DEGs decreased (D) or increased (E). (F,G) IPA diseases and functions enriched among decreased (F) or increased (G) DEGs (−log10 P values). (H,I) IPA upstream regulator analysis for decreased (H) or increased (I) DEGs. DEGs were defined by absolute fold change > 1.5 and FDR < 0.05 (n = 3 mice per group).

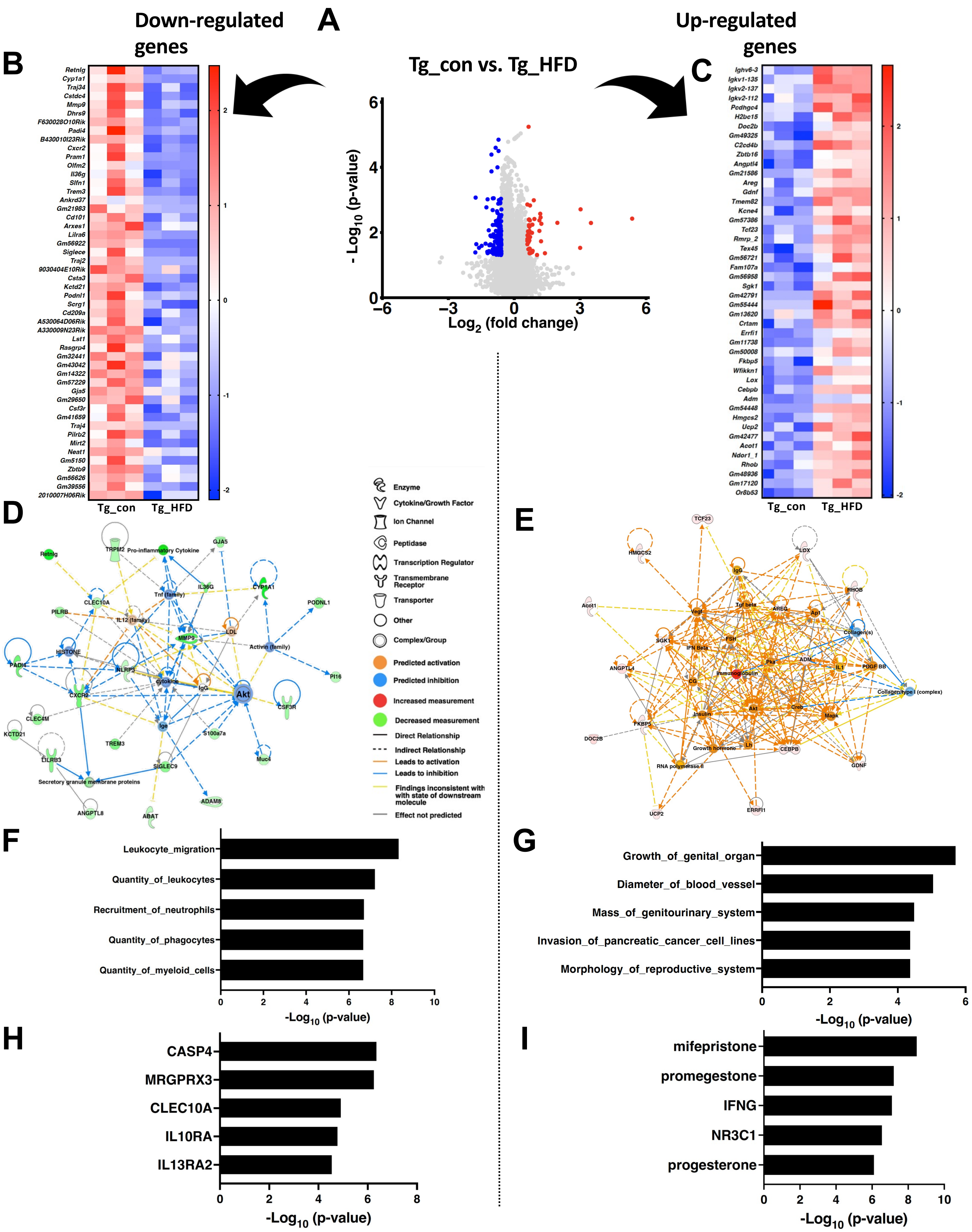


**Supplementary Figure S5. HFD induces transcriptional remodeling in βENaC-Tg lungs.** (A) Volcano plot of DEGs comparing Tg_con versus Tg_HFD. (B,C) Heatmaps of representative DEGs decreased (B) or increased (C) in Tg_HFD versus Tg_con (row z-scores). (D,E) IPA interaction networks generated from DEGs decreased (D) or increased (E). (F,G) IPA diseases and functions enriched among decreased (F) or increased (G) DEGs (−log10 P values). (H,I) IPA upstream regulator analysis for decreased (H) or increased (I) DEGs. DEGs were defined by absolute fold change > 1.5 and FDR < 0.05 (n = 3 mice per group).

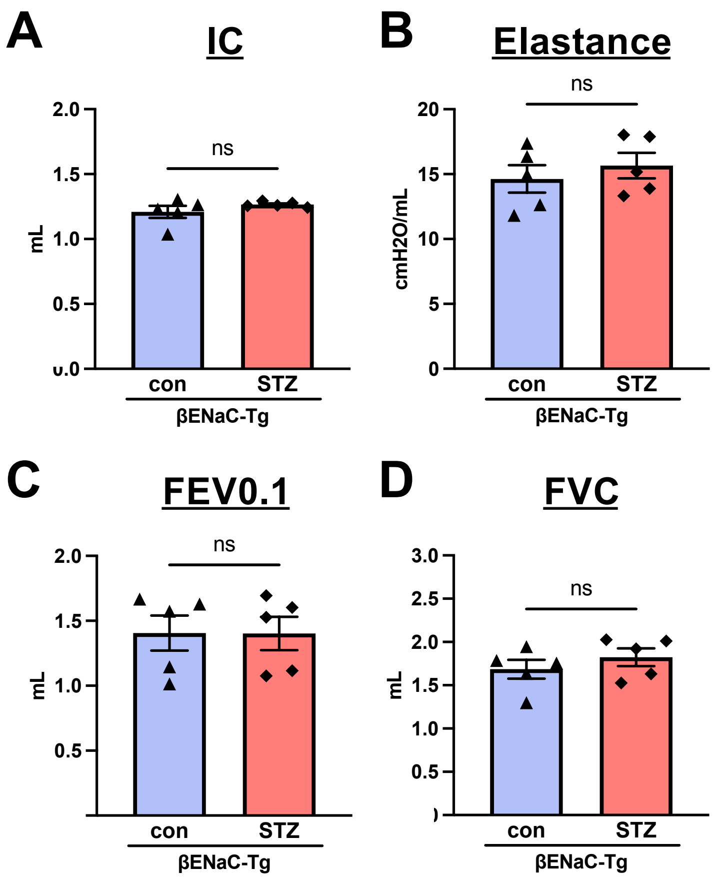


**Supplementary Figure S6. Additional pulmonary-function measurements after STZ treatment.** (A-D) Inspiratory capacity, elastance, FEV0.1, and FVC in vehicle- and STZ-treated βENaC-Tg mice. Data are mean ± SEM; each dot represents one mouse. n = 5-6 mice per group. Unpaired two-tailed Student t tests were used. ns, not significant.

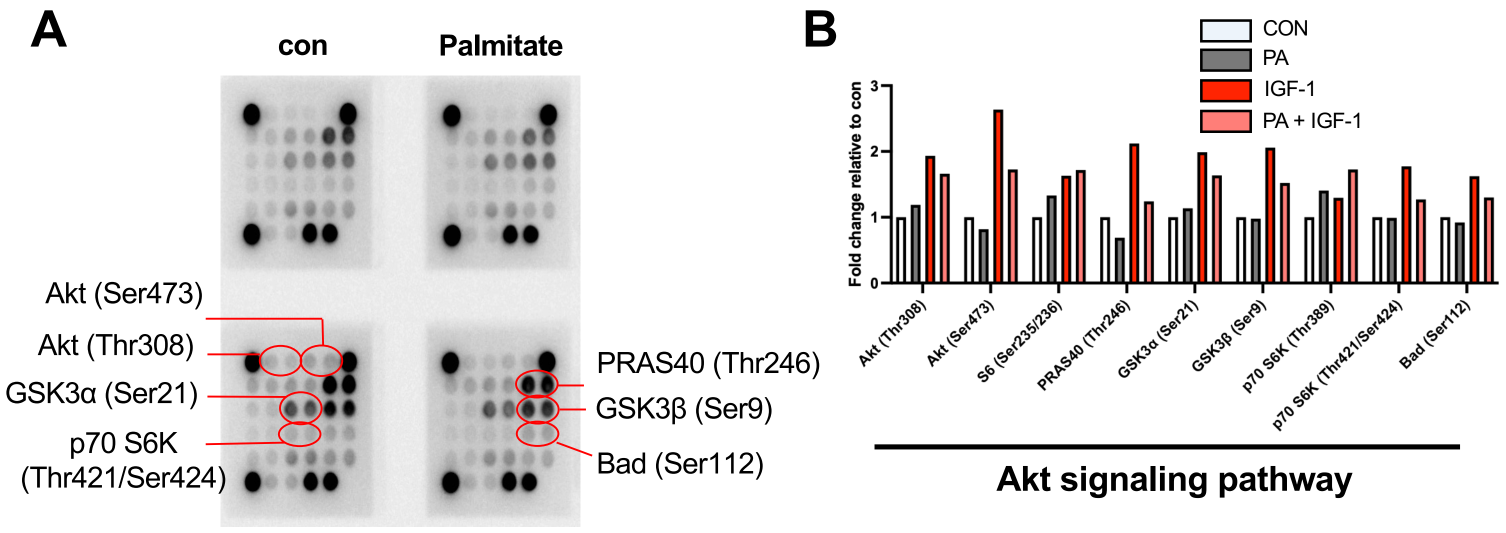


**Supplementary Figure S7. Palmitate broadly attenuates IGF-1-responsive Akt-pathway signals.** (A) PathScan Akt Signaling Antibody Array images from human bronchial epithelial 16HBE14o- cells exposed for 12 h to palmitate-free control medium (CON) or palmitate-containing medium (500 μM), followed by IGF-1 stimulation (50 ng/mL, 30 min). The four treatment conditions were CON, PA, CON + IGF-1, and PA + IGF-1. (B) Phosphorylation signals normalized to the array reference spots and expressed relative to CON. Signals from duplicate antibody spots were averaged. One array was analyzed per condition; therefore, the figure provides pathway profiling rather than inferential statistical analysis.

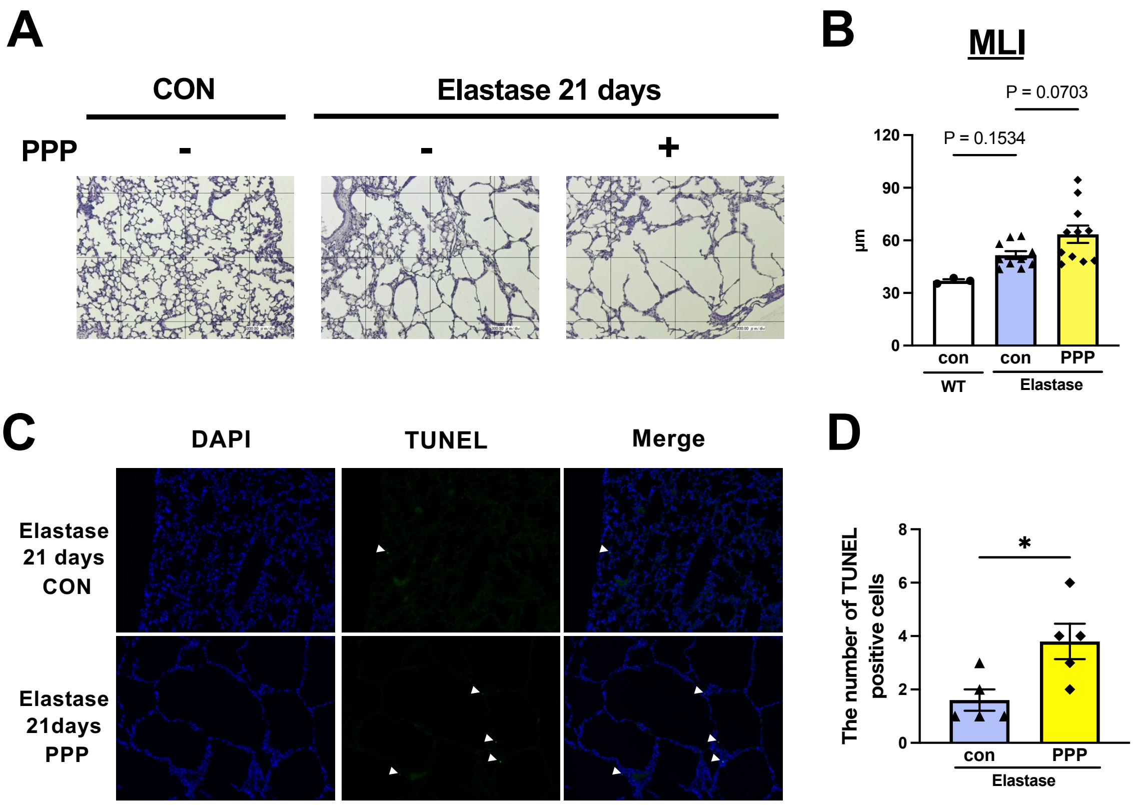


**Supplementary Figure S8. IGF-1R inhibition increases TUNEL positivity after elastase injury.** (A) Representative PAS and Alcian blue staining of airway mucus and bronchial morphology in elastase-injured mice treated with PPP (60 µg per day) for 2 weeks. (B) Quantitative morphometric analysis of alveolar septal architecture. (C) Representative TUNEL staining in elastase-injured mice treated with PPP. (D) Quantification of TUNEL-positive cells. Data are mean ± SEM; each dot represents one mouse (n = 5–11 mice per group). Statistics: one-way ANOVA with Dunnett's multiple comparisons. * $P < 0.05$.

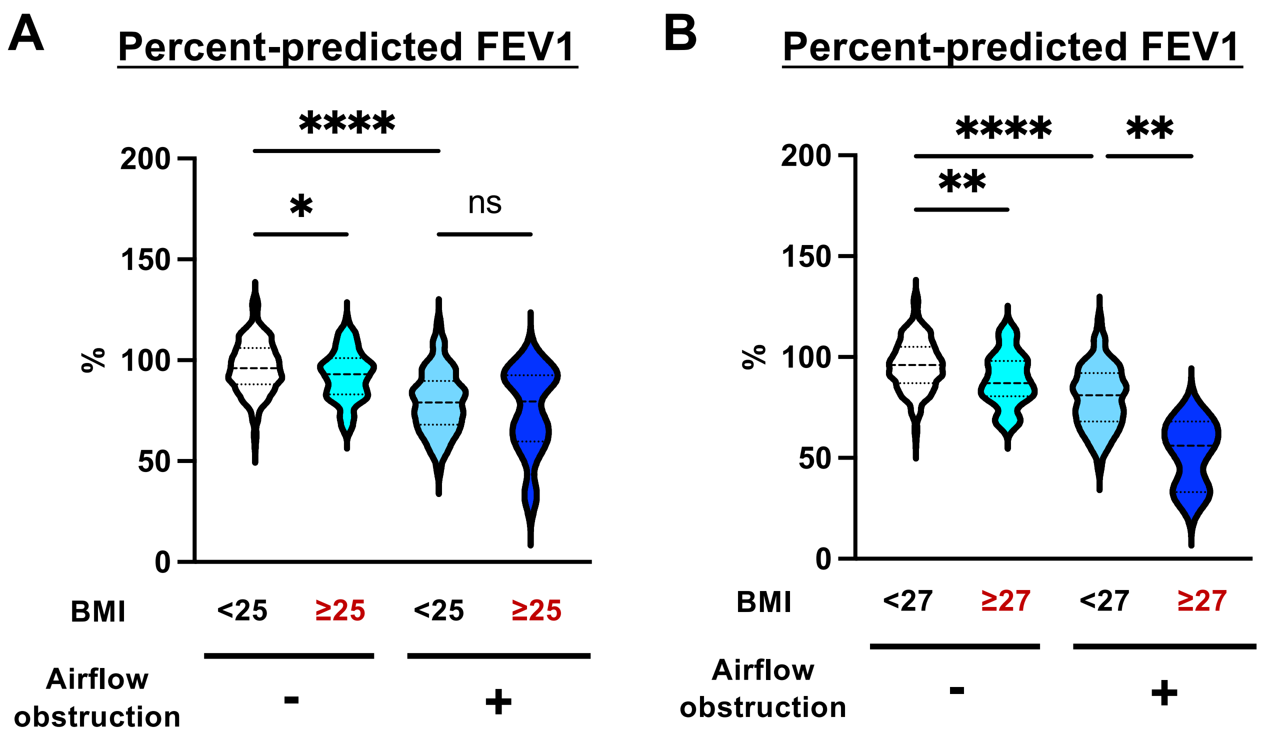


**Supplementary Figure S9. BMI-based sensitivity analyses of lung function according to airflow-obstruction status.** (A) Percent-predicted FEV1 stratified according to body mass index (BMI) and airflow-obstruction status using a BMI cutoff of 25 kg/m$^2$. From left to right, group sizes were n = 277, 121, 66, and 10. (B) Corresponding analysis using a BMI cutoff of 27 kg/m$^2$. From left to right, group sizes were n = 357, 41, 73, and 3. Airflow obstruction was defined as a pre-bronchodilator FEV1/FVC ratio <0.70. Violin plots show the distribution of individual values, and horizontal bars indicate medians. Because only three participants had both BMI ≥27 kg/m$^2$ and airflow obstruction, the analysis in (B) was considered exploratory. Statistical significance was assessed using one-way ANOVA followed by Tukey–Kramer multiple-comparison testing. ns, not significant; **P* <0.05; ***P* <0.01; *****P* <0.0001.
