## Supplementary material for "Dietary lipid stress aggravates obstructive lung injury involving epithelial IGF-1-Akt dysfunction and epithelial-endothelial crosstalk": uncropped blots

### Source data of Figure 2E: uncropped blots

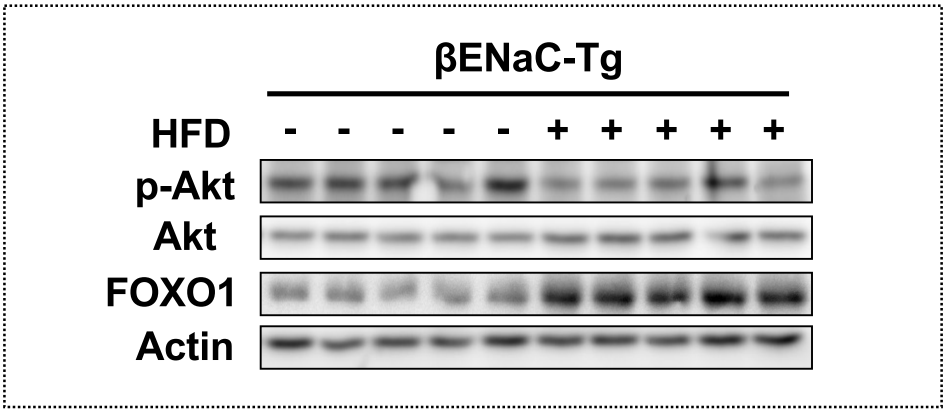


Akt

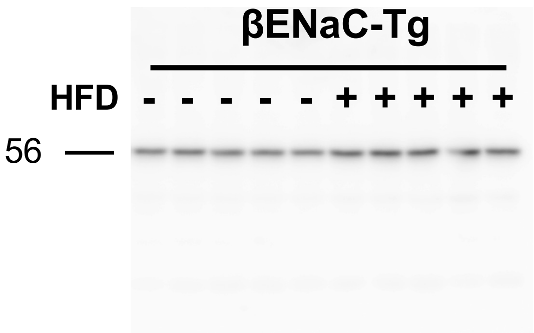


p-Akt

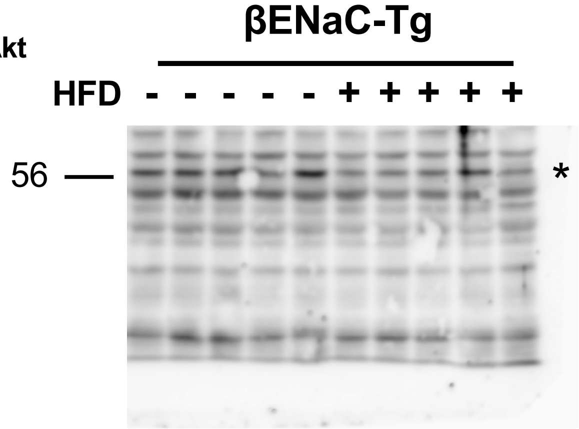


FOXO1

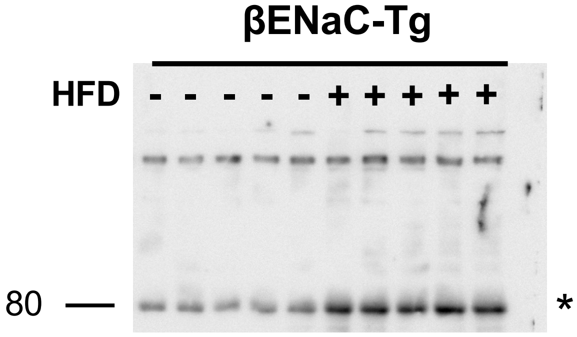


Actin

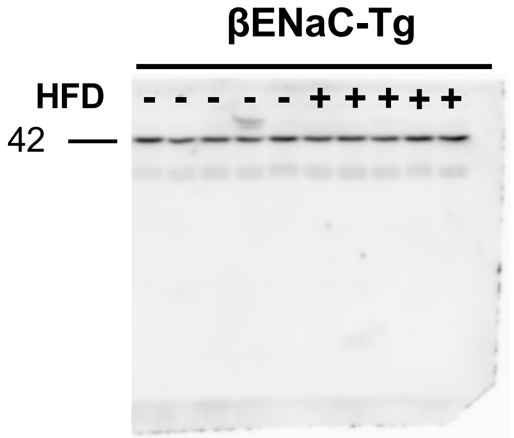


*Target bands

### Source data of Figure 2I: uncropped blots

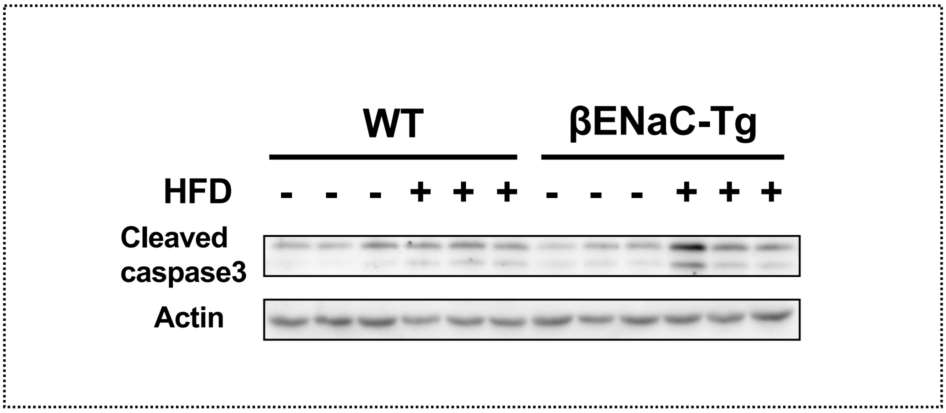


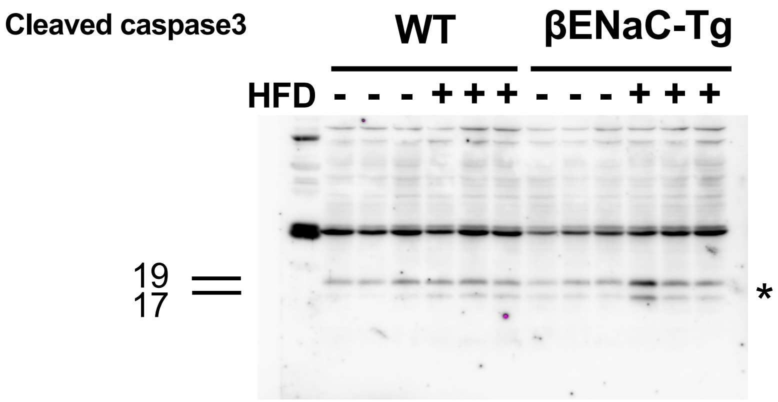


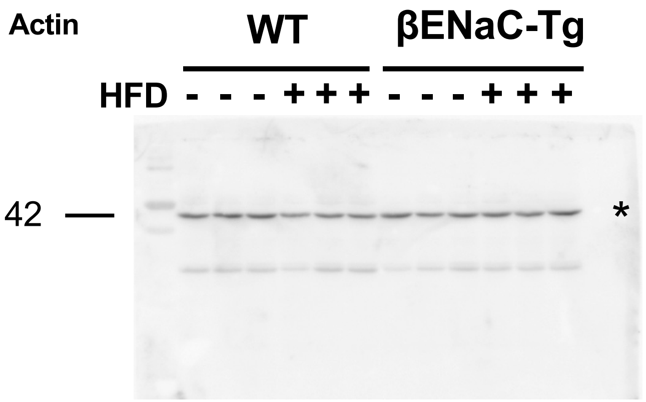


*Target bands

### Source data of Figure 3J: uncropped blots

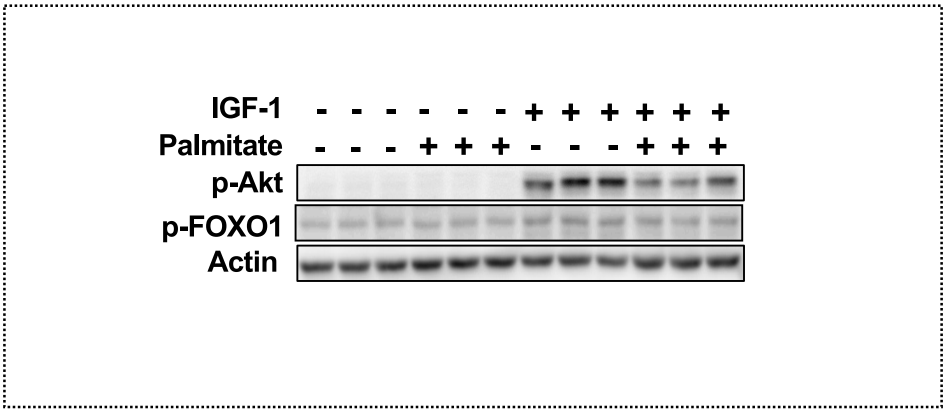


pAkt

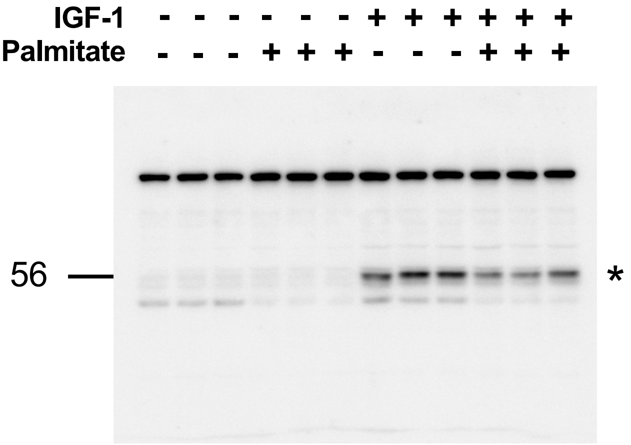


p-FOXO1

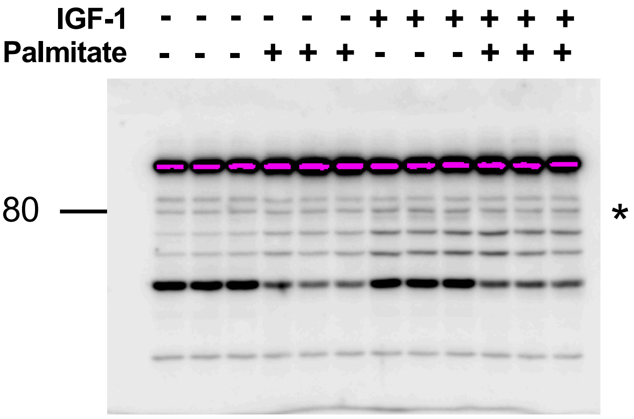


Actin

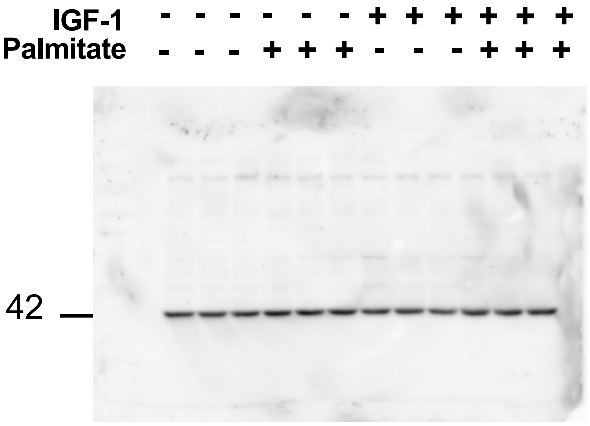


*Target bands

### Source data of Figure 4A: uncropped blots

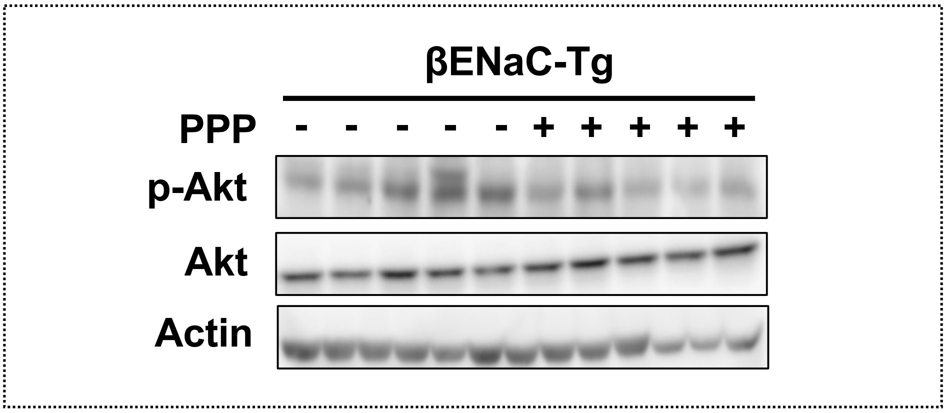


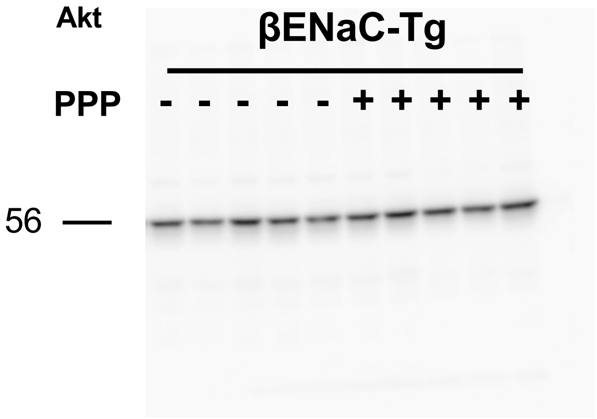


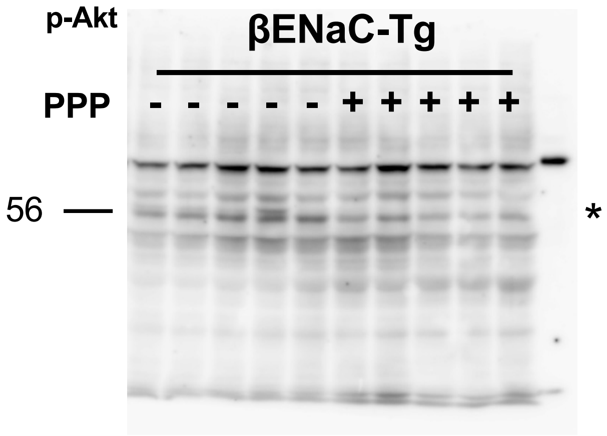


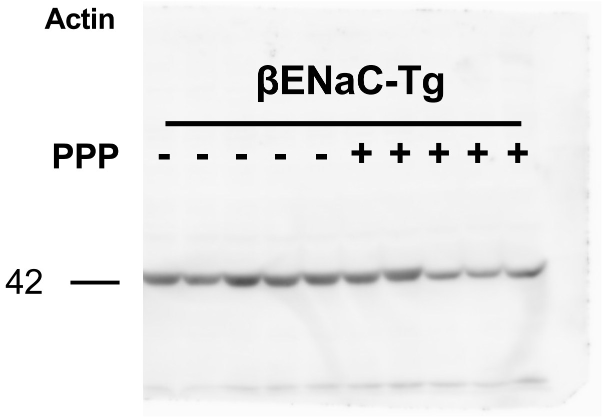


*Target bands

### Source data of Figure 5G: uncropped blots

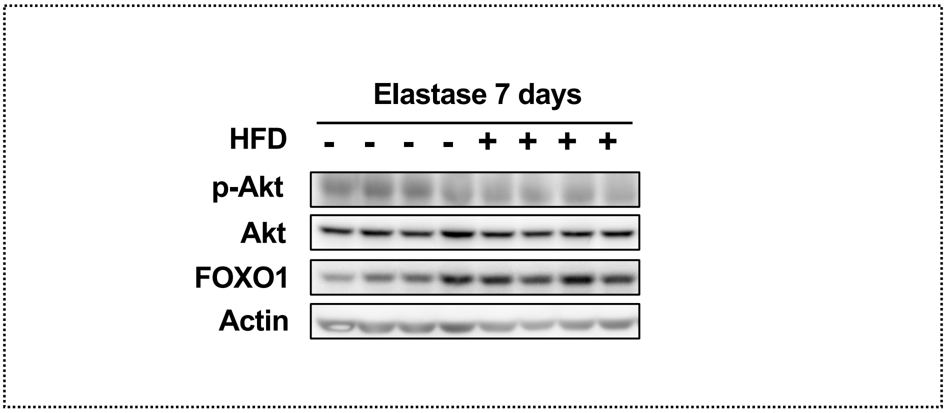


Akt

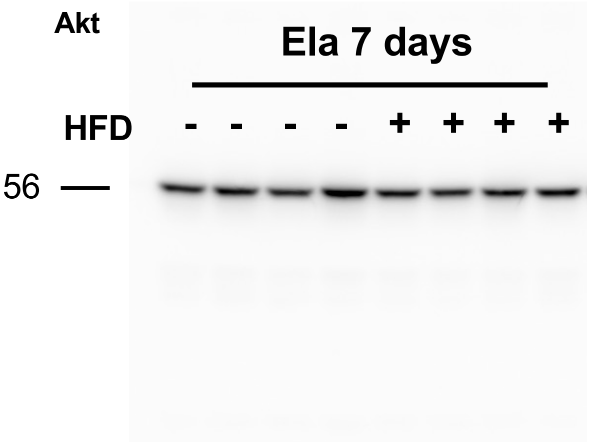


p-Akt

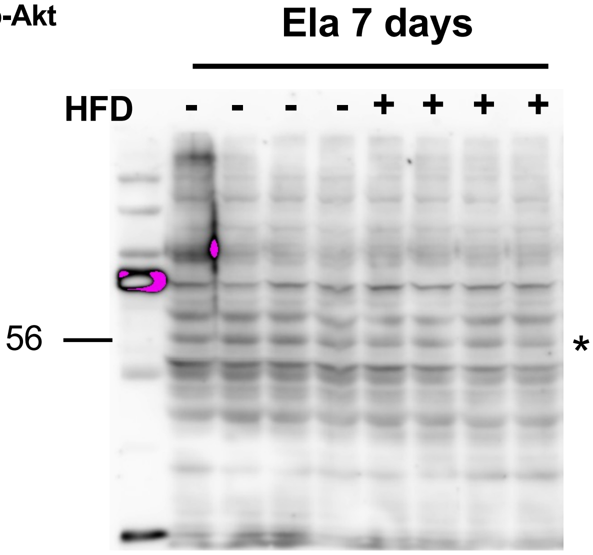


FOXO1

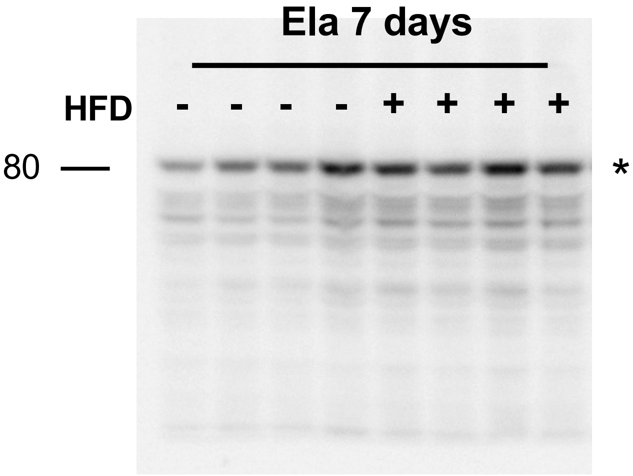


Actin

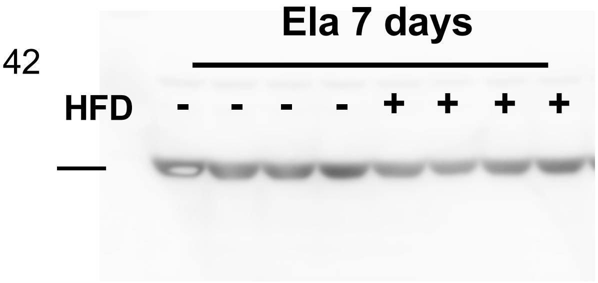


*Target bands

### Source data of Figure 6K: uncropped blots

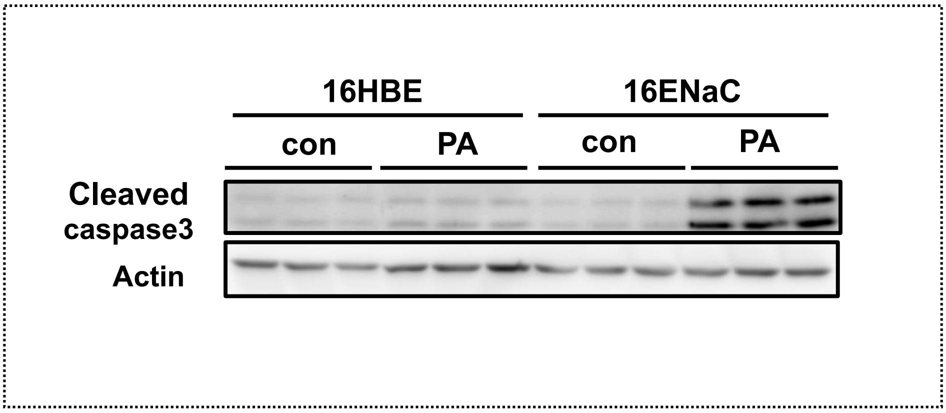


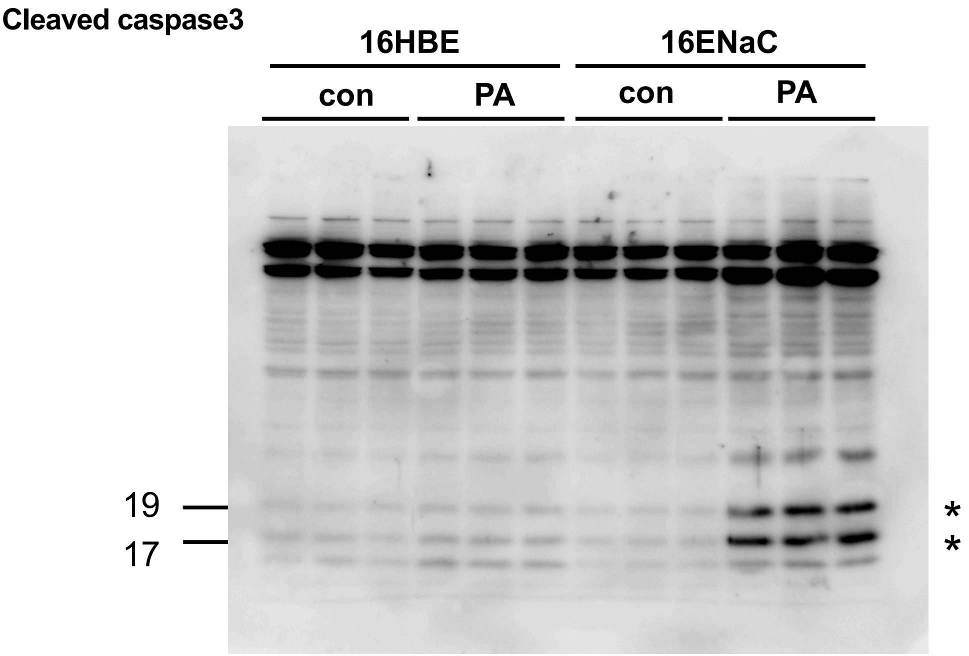


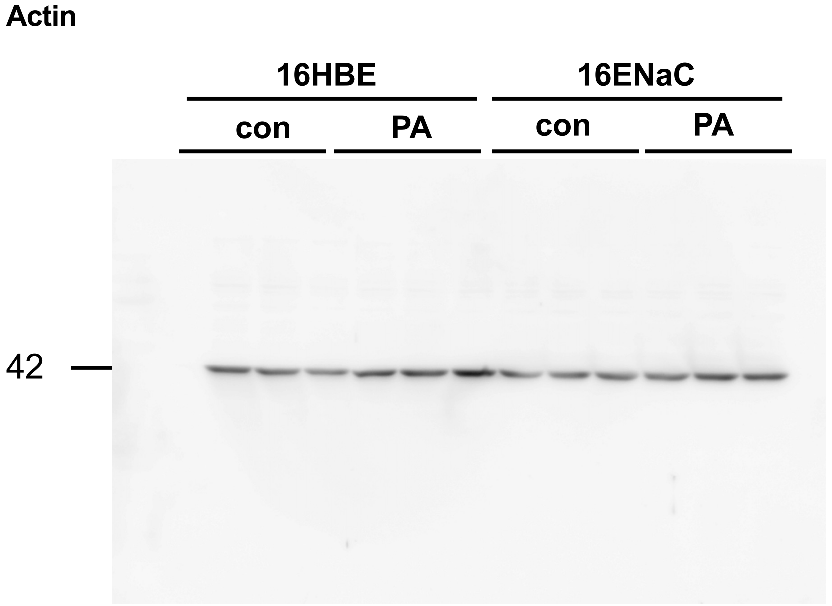


*Target bands
